# A cell death screen identifies macrophage-depleting agents with therapeutic potential

**DOI:** 10.64898/2026.09.17.752341

**Authors:** Stephan Forisch, Paul Ettel, Manuela Träger, Mario Mazic, Hannah K. Mayr, Andrea Vogel, Piyal Saha, Lovro Davidovski, Roko Sango, Velina S. Atanasova, Anna Gschwendtner, Christoph Trenk, Herwig P. Moll, Emilio Casanova, Helmut Dolznig, Markus Hengstschläger, Thomas Rattei, Anna Koren, Stefan Kubicek, Mario Mikula, Thomas Weichhart

**Author notes:** Contributed equally, listed in order of timely involvement (Stephan Forisch, Paul Ettel).

## Abstract

Macrophages are critical regulators of inflammation and tissue homeostasis, yet aberrant macrophage activation contributes to a wide spectrum of inflammatory and malignant diseases. Therapeutic strategies that directly reduce macrophage numbers have shown promise, but macrophage survival pathways remain incompletely defined, limiting the development of targeted therapeutic strategies. Here, we establish a high-throughput screening platform to identify small molecule inhibitors that impair macrophage survival. Screening a library of more than 2,000 targeted compounds in a cell survival assay, combined with *in silico* and *in vitro* analysis of preferential macrophage sensitivity, revealed the identification of three potent inhibitors: BIX-01294, GSK-J4, and Masitinib. All three compounds downregulated leukemia inhibitory factor receptor (LIFR), whose inhibition markedly reduced macrophage viability. *In vivo*, these inhibitors effectively depleted large peritoneal macrophages, ameliorated key symptoms of macrophage activation syndrome (MAS), and suppressed tumor growth in a syngeneic transplanted melanoma model as well as in an autochthonous lung cancer model. Together, these findings identify small molecule–mediated macrophage depletion as a promising therapeutic strategy and establish an experimental approach to uncover regulators of macrophage survival.

## Introduction

Macrophages are highly adaptable immune cells that contribute to tissue development, homeostasis, host defense, and repair. Their functions range from phagocytosis and metabolic support to orchestration of inflammatory and regenerative responses^1,2^. Macrophage functions are shaped by local cues, and they adopt diverse transcriptional and metabolic programs^1,3,4^. Under physiological conditions, macrophages sustain organ integrity and rapidly adjust to perturbations. Yet, in many diseases, macrophages adopt maladaptive activation states that amplify inflammation, drive tissue damage, or support tumor progression^1,5,6^.

Macrophage activation syndrome (MAS) represents a classical example where the pathological activity of macrophages is sufficient to cause disease. MAS, as potentially lethal disease, is considered as a type of secondary hemophagocytic lymphohistiocytosis (HLH) and can develop in patients suffering from the autoinflammatory disease systemic juvenile idiopathic arthritis (sJIA) or cancer^7^. Excessive stimulation by cytokines such as interferon-γ or granulocyte-macrophage colony stimulating factor (GM-CSF) leads to macrophage hyperactivation, hemophagocytosis, cytopenia, hepatosplenomegaly, and a severe cytokine storm^8,9^. While blockade of IL-1β or IL-6 provides partial protection, many patients remain at risk for MAS episodes, underlining the need for novel therapeutic strategies^10,11^.

Macrophages also play key roles in chronic inflammatory disorders and cancer. In many tumors, macrophages represent a large proportion of immune cells in the tumor microenvironment^12^ and contribute to tumor growth, immune suppression, angiogenesis, extracellular matrix remodeling, and therapy resistance^13–17^. These broad protumorigenic functions have stimulated intense interest in macrophage-directed therapies^12–14,16^.

Multiple strategies have been explored to modulate macrophage function, recruitment, or polarization in inflammatory diseases and cancer. Yet clinical success has been limited, fueling interest in approaches that directly deplete macrophages^18,19^. Although blockade of colony stimulating factor 1 receptor (CSF1R) can reduce macrophage numbers in preclinical models, the clinical potential of CSF1R inhibitors is constrained by compensatory signaling pathways, limited durability of responses, and genetic variants that reduce drug sensitivity^16,18^.

Together, these observations underscore the need for alternative therapeutic strategies that more effectively disrupt macrophage survival pathways. Small molecule inhibitors are particularly attractive due to their pharmacokinetics, tissue penetrance, and capacity to target intracellular processes essential for cell viability^20^. Yet, systematic efforts to identify macrophage-depleting compounds have been scarce, and unbiased discovery approaches remain underexplored.

Here, we present a high-throughput screening platform designed to identify small molecule inhibitors that preferentially impair macrophage survival. By integrating cell survival assays with *in silico* prioritization and *in vitro* cell-type sensitivity analyses, we identify three potent compounds with macrophage-depleting activity. We further demonstrate that these inhibitors reduce macrophage numbers *in vivo*, ameliorate key pathological features in a model of macrophage activation syndrome, and suppress tumor growth in both transplanted and autochthonous cancer models. These findings establish small molecule–mediated macrophage depletion as a promising therapeutic strategy for inflammatory and malignant diseases and provide a resource for future mechanistic interrogation of macrophage survival pathways.

## Methods

### Mouse Strains

C57BL/6J mice were bought from the core facility animal breeding and husbandry of the medical university of Vienna and from Janvier Labs, France. B6;129-Tsc2^fl/fl^ mice were crossed to Lyz2^cre/+^ mice to obtain *Tsc2^fl/fl^, Lyz2^cre/+^* mice, as described by Linke et al.^21^.

### Sex as a biological variable

Male mice used for the experiments. Mice were randomly assessed to groups and no blinding was used.

### Study approval

All mouse studies were approved by the official Austrian ethics committee for animal experiments (GZ 2023-0.380.996, GZ 2021-0.406.848). Mice were housed at a constant temperature and relative air humidity with a 12 h light/dark cycle. All experimental groups were age matched.

### *In vivo* Experimental Treatments

To investigate macrophage depletion, 8-week-old C57BL/6J mice were injected intraperitoneally (i.p.) with 20 mg/kg BIX-01294 (S8006, Selleckchem), 70 mg/kg GSK-J4 (S7070, Selleckchem), 30 mg/kg Masitinib (S1064, Selleckchem), or equal volumes of sterile PBS daily for one week. Alternatively, they received 300 µg anti-CD115 (C2169-50 mg, Leinco) or isotype control antibody (I-118-50 mg, Leinco) 3 times per week for 3 weeks via i.p. injection. For clodronate depletion, mice received 200 µl Clodronate Liposomes (Liposoma) or control liposomes (Liposoma) once daily for 2 days via i.p. injection.

For macrophage activation syndrome induction, 8-week-old C57BL/6J mice received 5 x 200 µg of the TLR9 agonist ODN 1826 (tlrl-1826, InvivoGen) *i.p.* as described before^22^. Two days after the first ODN 1826 daily treatments were performed as described above.

To generate a mouse melanoma model, 6-week-old C57BL/6J mice were intradermally injected with 5 x 10^5^ B16-F10 melanoma cells unilaterally. Two days after tumor induction, daily treatment as outlined above was started and tumor growth was measured daily. To measure tumor growth, following formular:

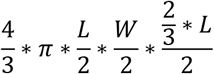

was used.

For the autochthonous lung cancer model, as previously described^23–25^ K-ras^LSL-G12D^ knock-in mice with p53 floxed alleles (K-ras^LSLG12D^:p53^fl/fl^; KP mice) were inhaled with 2.5 × 10^7^ plaque-forming units of a SPC-Cre recombinase–expressing adenovirus (Viral Vector Core, University of Iowa) and subsequently treated with the above describe dose of Masitinib.

### Cell Culture

Cells were cultured in a humidified CO_2_ (5 %) incubator at 37° C. For bone marrow isolation, femur and tibia were flushed with ice-cold DMEM (4.5 g/ L glucose, sodium pyruvate) containing 10 % fetal bovine serum (FBS, 26140079, Gibco), 10 % – 20 % L929 supernatant, 2 mM glutamine, 1X penicillin/ streptomycin (11548876, Gibco), 50 µg/ mL β-mercaptoethanol (11508916, Gibco) (Macrophage differentiation medium). Cells were either directly seeded on untreated 9 cm petri dishes or pelleted and resuspend in 100 µL FBS (26140079, Gibco) and 900 µL DMSO (5.89569, Sigma) for long-term storage in liquid nitrogen. Freshly seeded cells were split 1:2 after 3 days. Thawed cells had their media changed after 3 days and were split on day 4. After an additional 3 days, fully differentiated macrophages were harvested and seeded in DMEM (4.5 g/ L glucose, sodium pyruvate, 41965039, Gibco) containing 10 % fetal bovine serum, 20 ng/ mL recombinant murine M-CSF (315-02, PeproTech), 2 mM glutamine, 1 X penicillin/ streptomycin, 50 µg/ mL β-mercaptoethanol (Macrophage seeding medium) for downstream experiments. DLD-1, B16-F10, wild-type mouse embryonic fibroblasts (MEFs), and TSC1^-/-^ MEFs were cultured in DMEM (4.5 g/ L glucose, sodium pyruvate, 41965039, Gibco) containing 10 % FBS, 2 mM glutamine, 1 X penicillin/ streptomycin. Human umbilical vein endothelial cells (HUVECs) were cultured in endothelial cell growth medium 2 (C-22011, PromoCell) and passages three to five were used for experiments.

### Drugs

All drugs were solved in DMSO for *in vitro* experiments. Treatment times were as follows, 24 hours for dose-response curves, 8 or 16 hours for flow cytometric analysis, 16 hours for western blot analysis. For *in vivo* treatment regimes, all drugs were solved in distilled water (Gibco).

### High-Throughput Cell Viability Screen

Drugs part of the annotated compound library of the Chemical Screening Facility at the CeMM Molecular Discovery Platform were dispensed into 96-well plates for a final assay concentration of 10 µM using an acoustic liquid handler (Echo 550, BeckmanCoulter), and 2.5 x 10^4^ *Tsc2^fl/fl^, Lyz2^cre/+^* BMDMs were seeded on top using an automatic dispenser (MultiDrop Combi, ThermoFisher). Cells were cultured without the availability of M-CSF, as constitutive mTOR activation alone drives a hyper proliferative phenotype^21^. Cell viability was measured after 6 days using a CellTiter-Glo® assay (G7570, Promega) using an EnVision plate reader (Revvity), as per manufacturer’s instruction. Acquired signal readouts were normalized to the vehicle (DMSO) control.

### Cell Viability Assay (PrestoBlue™ Cell Viability Assay)

For metabolic cell viability analysis, cells were seeded in 96-well plates, treated with the indicated drugs and concentrations for 24 hours, before 11 µL PrestoBlue™ (A13261, Invitrogen) was added. After the corresponding incubation time, fluorescence intensity was measured using a Synergy HT Photometer. For DLD-1 and bone marrow-derived macrophages (BMDMs) 5 x 10^4^ and 2.5 x 10^4^ cells were seeded per well and incubated for 2 hours, respectively. For HUVECs, MEFs^WT^, and TSC1^-/-^ MEFs 4 x 10^3^, 3 x 10^3^, and 6 x 10^3^ cells were seeded per well and incubated for 1 hour, respectively.

### Cytotoxicity Assay (CyQUANT™ LDH Cytotoxicity Assay)

To measure LDH release a CyQUANT™ LDH cytotoxicity assay (C20300, ThermoFisher) was used as per manufacturers instruction. Briefly, BMDMs were seeded at 2.5 x 10^4^ cells per well in a tissue culture treated 96-well plate. After treatment with the indicated drugs, concentrations, and time, 50 µL of supernatant was transferred to a new 96-well plate. To each sample well, 50 µL reaction mixture was added and the plate incubated for 30 minutes at room temperature. After this, 50 µL of stop solution was added to each sample well. Absorbance was measured at 490 nm and 680 nm using a Synergy HT photometer (Biotek). Acquired readings were normalized to the background and maximum LDH release wells, as per manufacturer’s instructions.

### mRNA isolation, library preparation and bulk RNA sequencing

BMDMs were treated with BIX-01294 (24h, 1.85 µM), GSK-J4 (4h, 12.3 µM), or Masitinib (16h, 8.5 µM). RNA was extracted using Monarch® RNA Cleanup Kit (New England Biolab®, T2040). The amount of total RNA was quantified using the Qubit 2.0 Fluorometric Quantitation system (Thermo Fisher Scientific, Waltham, MA, USA) and the RNA integrity number (RIN) was determined using the 2100 Bioanalyzer instrument (Agilent, Santa Clara, CA, USA). RNA-seq libraries were prepared with the NEBNext® Ultra™ II Directional RNA sample preparation kit (New England Biolabs, Inc., Ipswich, MA, USA). NGS library concentrations were quantified with the Qubit 2.0 Fluorometric Quantitation system (Life Technologies, Carlsbad, CA, USA) and the size distribution was assessed using the 2100 Bioanalyzer instrument (Agilent, Santa Clara, CA, USA). For sequencing, samples were diluted and pooled into multiplex NGS libraries in equimolar amounts.

NGS reads were mapped to the Genome Reference Consortium GRCm38 assembly via “Spliced Transcripts Alignment to a Reference” (STAR, 2.7.9a) utilizing the “basic” Ensembl transcript annotation from version e100 (April 2020) as reference transcriptome. Since the mm10 assembly flavor of the UCSC Genome Browser was preferred for downstream data processing with Bioconductor packages for entirely technical reasons, Ensembl transcript annotation had to be adjusted to UCSC Genome Browser sequence region names. STAR was run with options recommended by the ENCODE project. NGS read alignments overlapping Ensembl exon features were counted with the Bioconductor (3.14) GenomicAlignments (1.30.0) package via the summarizeOverlaps function in Union mode, ignoring secondary alignments and alignments not passing vendor quality filtering. Since dUTP-based RNA-seq protocols lead to the sequencing of the first strand, all alignments needed inverting before strand-specific counting in feature (i.e., gene, transcript, and exon) orientation. Exon-level counts were aggregated to gene-level counts and the Bioconductor DESeq2 (1.34.0) package was used to test for differential expression based on a model using the negative binomial distribution.

An initial exploratory analysis included principal component analysis (PCA), multi-dimensional scaling (MDS) and sample distance heatmap plots (ggplot2, 3.3.6), all annotated with variables used in the expression modelling.

Biologically meaningful contrasts were extracted from the model, log2-fold values were shrunk with the CRAN ashr (2.2.-54) package, while two-tailed p-values obtained from Wald testing were adjusted with the Bioconductor Independent Hypothesis Weighting (IHW, 1.22.0) package. The contrasts were visualized via MA, volcano (Bioconductor EnhancedVolcano, 1.12.0) and expression heatmap (Bioconductor ComplexHeatmap, 2.10.0) plots. The resulting gene tables were annotated and subsequently filtered for significantly differentially up- and down-regulated genes, and independently subjected to gene set enrichment analysis (Enrichr).

### Peritoneal Macrophage Isolation

Mice were euthanized by ketamine (Ketasol, Livisto) and xylazine (Rompun®, Bayer) overdose. The skin covering the abdomen was cut and pulled to the side to expose the peritoneal cavity. A 27G needle was used to inject 5 mL FACS buffer (DPBS without Ca2+, Mg2+ containing 2 % FBS and 2.5 mM EDTA, all Gibco) into the peritoneal cavity. After gently massaging the abdomen, peritoneal exudate was removed using a 25G needle. The procedure was repeated once. Isolated peritoneal exudate was stored on ice for further downstream processes.

### Flow Cytometry

Differentiated BMDMs were harvested and seeded in laminar wash plates. After drug treatment for 8 or 16 hours (EC359, HY-120142, MCE) with the indicated concentrations, wells were washed with Annexin V binding buffer (DPBS without Ca^2+^, Mg^2+^ containing 2 % FBS and 2.5 mM CaCl_2_) using a HT-2000 laminar wash system (Curiox). Following a 20-minute incubation with Annexin V (2.5 µL, 640905, BioLegend) and 7-ADD (1 µg/ mL, SML1633, Sigma), wells were acquired using a CytoFLEX S (Beckmann Coulter). Peritoneal exudate was spun down, resuspended in 1 X red blood cell lysis buffer (420301, BioLegend) and incubated for 10 minutes on ice. Pellets were washed with FACS buffer. Equal number of cells were blocked using TruStain FcX (10131950, BioLegend) for 10 minutes and stained with the indicated antibodies (all at 1:200 dilutions) for 30 minutes, both incubations were performed on ice and in the dark. Pellets were washed with FACS buffer and resuspended in PBS containing 7-AAD. After 30 minutes of incubation in the dark on ice, cells were directly acquired.

### Histology

Mouse tissues were fixed overnight in Histofix (P087.1, Roth), washed with DPBS, and processed using a KOS microwave tissue processor before embedding in paraffin for long term preservation. All tissues were cut at a thickness of 2 µM using a microtome (Leica). Formaldehyde fixed, paraffin embedded (FFPE) sections were cleared using neoclear (1.09843.5000, Merck) and antigenic epitopes were retrieved by heating slides immersed in citrate buffer (pH 6.0, S236984-2, Dako) for 10 minutes at 120°C using an autoclave. These sections were further processed in downstream applications described below. For Hematoxylin & Eosin staining slides were stained with Mayer’s Hematoxylin and Eosin Y and mounted with Neomount.

### Immunohistochemistry (IHC)

Endogenous peroxidases were blocked by incubating slides with 3 % H_2_O_2_ (H1009, Sigma) for 10 minutes. Endogenous biotin and streptavidin were blocked by incubating with the respective blocking solutions (SP-2001, VectorLabs) for 15 minutes each. Unspecific epitopes were blocked by incubation with PBS-T containing 2.5 % goat serum for 20 minutes. Sections were then incubated with PBS-T containing either an anti-Mac-2 (1:10000, CL8942AP, Cedarlane) or F4/80 (1:400, 70076, Cell Signaling) antibody overnight at 4°C. On the next day, biotinylated goat anti-rat or goat anti-rabbit IgG antibodies (1:500) were added to sections and incubated for 45 minutes. Afterwards, sections were incubated with Novocastra streptavidin conjugated HRP (RE7110-CE, Leica) for 30 minutes. AEC high sensitivity chromogen substrate (Code K3461, Dako) was used for detection. Mac-2-stained slides were incubated for 5 minutes. For nuclear counterstaining, slides were immersed in filtered haematoxylin for 10 seconds. Slides were washed to remove excess hematoxylin (1051740500, Merck) and mounted using Aquatex® (1.08562.0050, Merck). Whole-mount tissue sections were acquired using an Olympus BX63 microscope (Olympus).

### *In-Situ* detection of Apoptosis

For *in-situ* detection of late apoptotic cells, a Click-iT™ Plus TUNEL Assay (Alexa Fluor 647, C10619, Invitrogen) was used. Proteinase K treatment was omitted as it degraded the CD45 epitope. Sections were incubated with TdT reaction buffer for 10 minutes at 37°C before adding the TdT reaction mix and incubating for 60 minutes at 37°C. After rinsing of slides, sections were incubated with TUNEL reaction cocktail for 30 minutes at 37°C. Following completion of the TUNEL assay, sections were blocked using PBS-T containing 2.5 % donkey serum for 20 minutes. Antibodies against CD45 (1:100, 70257, Cell Signaling) and F4/80 (1:100, 123102, BioLegend) were diluted in PBS-T and sections incubated overnight at 4°C. On the next day, donkey anti-rat Alexa Fluor 488 (1:500, Invitrogen) and donkey anti-rabbit Alexa Fluor 555 (1:500, Invitrogen) were diluted in PBS-T and sections incubated for 45 minutes. After completion of immunofluorescent staining, autofluorescence was quenched using by incubation with Vector TrueVIEW (VectorLabs) reagent for 5 minutes. Slides were mounted with VECTASHIELD Vibrance antifade mounting media (VectorLabs) and images acquired using an Olympus BX63 (Olympus) epifluorescent microscope.

### Immunofluorescent Staining

Unspecific antibody binding was blocked using 2.5% serum diluted in PBS with 0.025% Triton X (PBST). Sections were incubated with primary antibodies (Rat anti-F4/80 (1:100, 123102, BioLegend), Rabbit anti-CD45 (1:200, 70257, Cell Signaling), Goat anti-CD206 (1:100, PA5-46994, Invitrogen), Rabbit anti-pSTAT1 (1:50, 9171, Cell Signaling), Rabbit anti-CD31 (1:100, 77699, Cell Signaling Technology), Rabbit anti-CD8 (1:200, 0648R, Bioss), Rabbit anti-SOX10 (1:100, ab180862, abcam), Rat anti-Ki-67 (1:100, 14-5698-82, ThermoFisher Scientific) in PBST and incubated overnight. The following day, sections were incubated with secondary antibodies and DAPI for 45 min. and mounted using Fluoromount™ (F4680, Sigma). Images were acquired using an Olympus BX63 (Olympus) epifluorescent microscope.

### Western Blotting

Cells were seeded in untreated petri dishes or untreated 6 or 12-well plates. After drug treatment for the indicated concentration and time, media was aspirated, and wells washed twice with ice-cold DPBS. Cells were scraped in 1 X RIPA buffer (ab156034, abcam) supplemented with cOmplete™ (11697498001, Roche) and PhosSTOP protease and phosphatase inhibitors (PHOSS-RO, Roche), 20 µg/ mL trypsin inhibitor (T9003, Sigma), 4 µg/ mL aprotinin (A1153, Sigma), 4 µg/ mL leupeptin (L2884, Sigma), 0.6 µg/ mL benzamidinchlorid (8.20122, Sigma), and 2 mM AEBSF (A8456, Sigma). To remove cellular debris and DNA samples were spun down for 15 minutes at 12.000 rpm and at 4°C. Protein concentration was measured using a Pierce™ Rapid Gold BCA assay (A53226, Invitrogen). Protein lysates were boiled at 65°C for 10 minutes in protein sample loading buffer containing 100 mM DTT (10197777001, Sigma). Equal amounts of protein lysates were separated using a 13.5 % SDS-PAGE (L3771, Sigma) system and transferred to nitrocellulose membranes (P/N 926-31092, LI-COR) using either a Trans-Blot Turbo transfer system or Mini Trans-Blot Cell (both Bio-Rad). Subsequently, membranes were blocked in 0.5 X Intercept (TBS) blocking buffer (927-60001, LI-COR) for 1 hour at room temperature. Antibodies (Rabbit anti-pRIPK3 (1:1000, 91702, Cell Signaling), Rabbit anti-cleaved-caspase 3 (1:500, 9664S, Cell Signaling), Rabbit anti-GSDMD (1:1000, ab219800, abcam), Rabbit anti-cleaved-PARP (1:1000, 9532T, Cell Signaling), Mouse anti-β-actin (1:2500, A1978, Sigma)) were diluted in 0.5 X Intercept (TBS) blocking buffer (927-60001, LI-COR) containing 0.1 % Tween-20. Membranes were incubated with diluted antibody solutions overnight at 4°C on a tube roller. Secondary goat anti-mouse 680 RD and goat anti-rabbit 800 CW antibodies (both 1:20000) were diluted in 1 X Intercept (TBS) Antibody Diluent and membranes incubated for 1 hour at room temperature. Membranes were scanned using the auto sensitivity setting of a LI-COR Odyssey cLX (LI-COR).

### Image Analysis

Whole mount IHC and IF images were analyzed using modules available in HALO (Indica Labs) and Strataquest (Tissuegnostics). Data was normalized as indicated.

### Publicly available RNA sequencing analysis

Microarray or RNA sequencing from isolated monocytes from sJIA were downloaded from the Gene Expression Omnibus (GEO) database (GSE80060 and GSE147608) and analyzed for differential expression of the target genes. Bulk RNA sequencing from melanoma patients was downloaded from the University of California Santa Crut Xena Functional Genomics Explorer (TCGA TARGET GTEx data set). Correlations between the genes of interest were calculated using the linear regression function in GraphPad Prism (Version 9.0.2).

### Statistical Analysis

Data was graphed and analyzed with GraphPad Prism (Version 8.3). Groups were compared using an unpaired two-tailed t-test or one-way ANOVA depending on the number of groups. Statistical significance is indicated for *p ≤ 0.05*, *p ≤ 0.01*, *p ≤ 0.001*, *p ≤ 0.0001* with *\**, *\*\**, *\*\*\**, *\*\*\*\**, respectively.

### Data and materials availability

Bulk RNA-seq data generated in this study have been deposited at GEO and are publicly available (GSE268965). This paper analyzes existing, publicly available RNA-seq data (GSE80060 and GSE147608).

## Results

### A high throughput screen identifies drugs with macrophage depleting potential

To discover compounds that interfere with macrophage survival pathways, we performed a high-throughput viability screen using bone marrow–derived macrophages (BMDMs) generated from *Tsc2^fl/fl^, Lyz2^cre/+^*mice^21^. As previously reported, these macrophages display constitutive mTORC1 activation and increased proliferative capacity^26–28^. Furthermore, they survive without exogenous CSF1, thereby enabling the identification of growth factor-independent survival pathways. We screened 2,171 targeted small molecules with defined molecular targets, primarily acting on receptor tyrosine kinases or epigenetic regulators, at a concentration of 10 μM^29^ (Figure 1A). After six days of treatment, macrophage viability was quantified using a luminescent ATP-based assay following established high-throughput screening protocols^29^. From this screen, we identified 228 compounds that reduced macrophage viability by at least 60% relative to DMSO controls (Figure 1B). These compounds were selected for further analysis. To identify pathways associated with the observed loss of viability, we performed bioinformatic enrichment using Enrichr^30^ and Drugmonizme^31^ (Figure 1A). As expected, compounds targeting the PI3K-Akt-mTOR were enriched, consistent with the constitutive activation of mTORC1 in *Tsc2^fl/fl^, Lyz2^cre/+^*macrophages. Target annotation further highlighted the involvement of receptor tyrosine kinases (RTKs) as well as chromatin-modifying enzymes and cell cycle regulators (Figures 1C, S1A-C).

**Figure 1:**
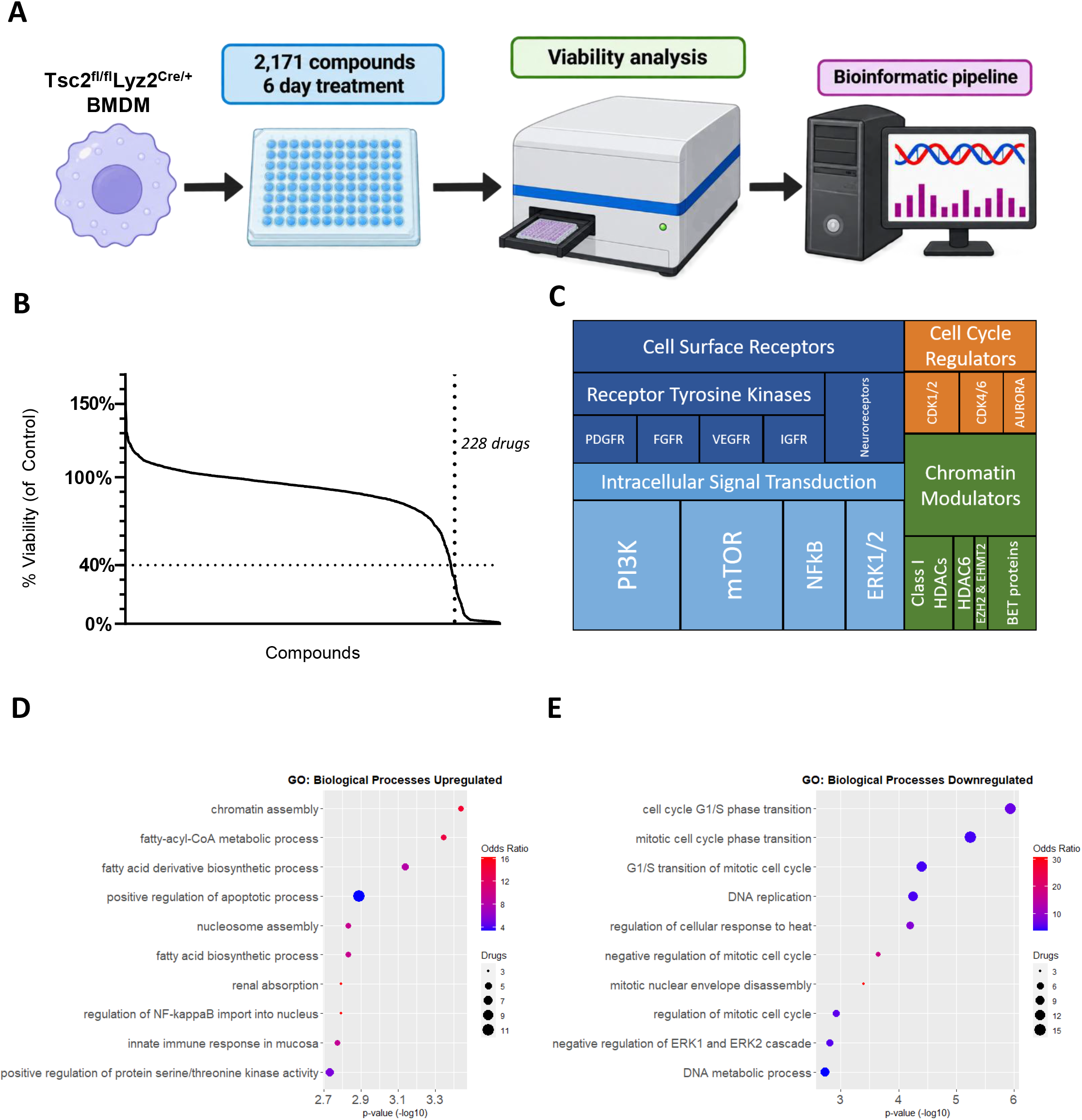
Identification of compounds affecting macrophage viability. (**A**) Schematic overview of the high-throughput screening workflow used to identify compounds that reduce macrophage viability. (**B**) Ranked distribution of compounds tested in the screen based on their effect on macrophage viability. The dotted line indicates the threshold corresponding to a 40 percent reduction in viability. (**C**) Tree flow diagram illustrating the frequency of annotated molecular targets among compounds that reduced macrophage viability in the screen. (**D**, **E**) Gene ontology analysis of biological pathways enriched among targets of compounds that decreased viability of Tsc2^fl/fl^, Lyz2^cre/+^ bone marrow-derived macrophages in the high-throughput screen, showing pathways associated with upregulated (D) and downregulated (E) targets.

Gene ontology analysis of the drug targets also revealed enrichment in fatty acid metabolism genes (Figure 1D) consistent with studies linking metabolic stress to cell death^32^. Previous work showed that the hyperproliferative phenotype of *Tsc2^fl/fl^, Lyz2^cre/+^*BMDMs is associated with upregulation of mitotic cell cycle transition pathways^21^. Consistent with this, these pathways were downregulated in response to the inhibitory compounds in our screen (Figure 1E). Together, these findings identify a set of compounds that reduce macrophage viability and highlight the roles of RTK signaling, chromatin regulation, metabolic pathways, and cell cycle control in supporting macrophage survival.

### Selected compounds induce cell death of macrophages

To identify compounds with preferential toxicity toward macrophages while sparing other cell types, we queried the 228 hits from the primary screen in PharmacoDB^33^ to obtain IC_50_ values across a large panel of cancer and non-cancer cell lines (Figure 2A). Compounds were ranked based on the percentage of cell lines with IC_50_ values above 10 μM and the top 15 drugs were selected for further *in vitro* evaluation (Figure 2B). This subset retained the broad pathway diversity observed in the initial screening results. For subsequent experiments, TSC2^fl/fl^ BMDMs were included as controls to evaluate compound activity independent of chronic mTORC1 signaling. To confirm the screening results, we first measured lactate dehydrogenase (LDH) release as a marker of membrane lysis. In control *Tsc2^fl/fl^* BMDMS, most compounds induced significantly increased LDH release only after 24 hours, but not after 12 hours (Figure 2C), indicating that they triggered a slow form of cell death. Further Annexin V and 7-AAD staining indicated that the majority of the identified drugs except GSK-J4 retained the cells largely intact (Annexin V^+^, 7-AAD^-^) indicating apoptosis rather than early membrane permeabilization (Figure 2D). Based on LDH kinetics, an 8-hour treatment window was used to avoid secondary necrosis, which becomes prominent after 12 hours in *Tsc2^fl/fl^, Lyz2^cre/+^* BMDMs.

**Figure 2:**
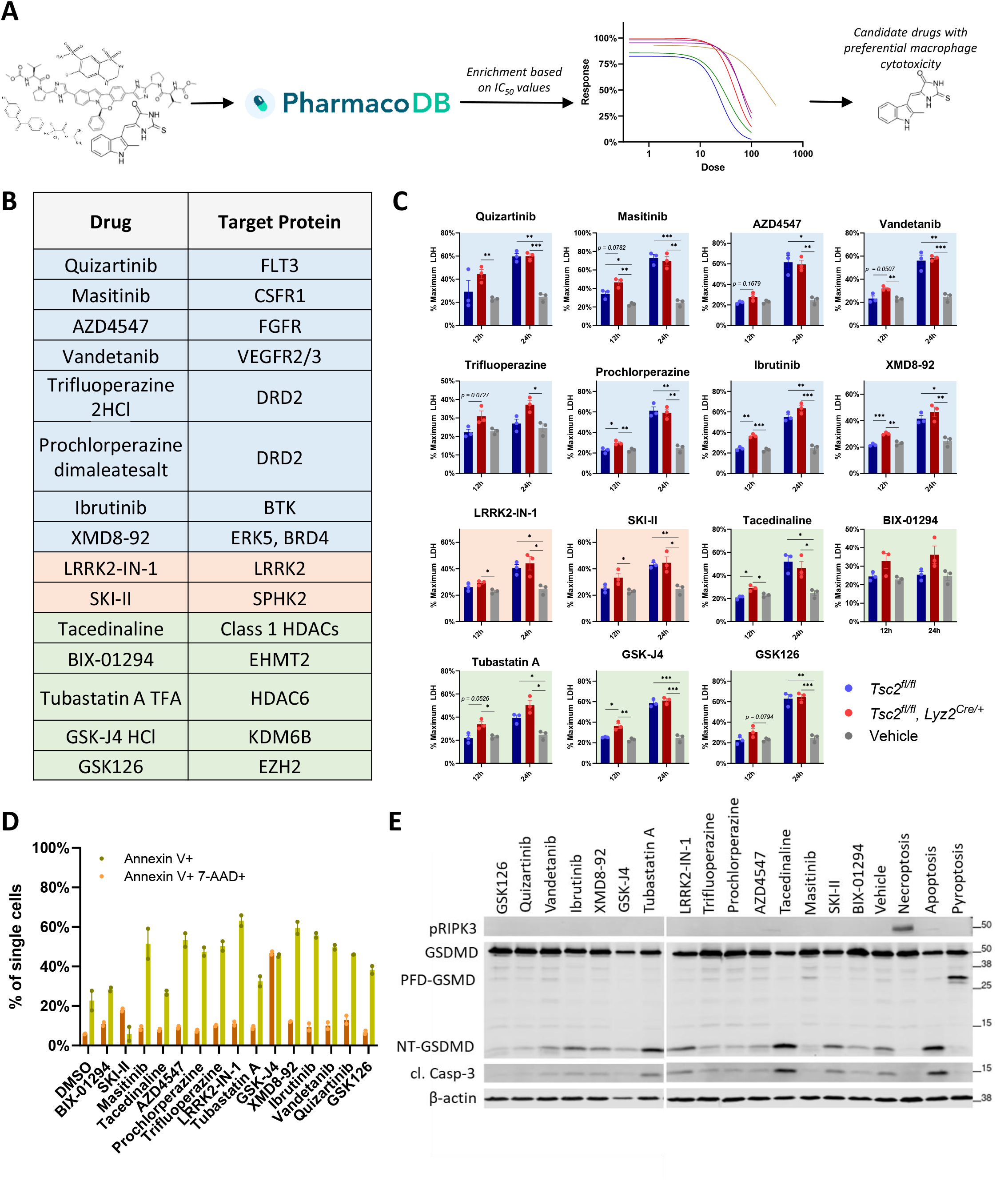
Selected compounds induce cell death in macrophages. **(A)** Workflow illustrating the prioritization and enrichment of candidate compounds with preferential macrophage cytotoxicity. **(B)** Candidate compounds identified from the screen and their primary molecular targets. Receptor tyrosine kinases (RTKs) and downstream signaling components are shown in blue, chromatin-modifying enzymes in green, and other cytoplasmic kinases in rosé. IC50 values (Table 2) were used for subsequent assays to assess the mode of cell death induced by each compound, including LDH release **(C)**, Annexin V/7-AAD staining **(D)**, and immunoblot analysis **(E)**. **(C)** LDH release from Tsc2^fl/fl^ and Tsc2^fl/fl^, Lyz2^cre/+^ bone marrow-derived macrophages treated with candidate compounds for 12 or 24 hours. Background shading corresponds to compound classes shown in **(B**). Data represent mean ± SEM of three independent experiments with three technical replicates each. Vehicle controls were intentionally shared across compounds tested within the same experimental batch, as these drugs were assessed simultaneously on the same plates using the same control wells. **(D)** Flow cytometric analysis of apoptosis and necrosis in C57BL/6J bone marrow-derived macrophages treated for 8 hours with IC50 concentrations of the indicated compounds. Data represent mean ± SEM of two technical replicates. **(E)** Immunoblot analysis of C57BL/6J bone marrow-derived macrophages treated for 16 hours with the indicated compounds. Apoptosis was induced by incubation with 1 µM staurosporine for 4 hours. Pyroptosis was induced by stimulation with 100 ng/mL LPS for 2 hours followed by 10 µM nigericin for 30 minutes. Necroptosis was induced by treatment with 100 ng/mL LPS and 50 µM Z-VAD-FMK for 2 hours. Statistical significance is indicated as * p < 0.05, ** p < 0.01, *** p < 0.001.

To determine whether pyroptosis or necroptosis contributed to cell death, we analyzed key markers of these pathways. For pyroptosis, we assessed cleavage of GSDMD, which generates either the pore-forming PFD-GSDMD fragment via caspase-1/11 or the smaller, non-lytic NT-GSDMD fragment via caspase-3^34^. Several compounds induced NT-GSDMD, but no PFD-GSDMD was detected (Figure 2E). For necroptosis, we tested for phosphorylated RIPK3 (pRIPK3), the hallmark of necrosome activation^35^, but no pRIPK3 was detected (Figure 2E). Collectively, these data indicate that the selected compounds initially trigger predominantly apoptosis-like cell death, followed by membrane permeabilization at later time points, particularly in Tsc2-deficient macrophages, without evidence for pyroptosis or necroptosis as dominant cell-death pathways.

### Three drugs show preferential macrophage killing potential *in vitro* and modulate LIFR expression

To assess whether the selected compounds preferentially affect macrophages, we generated dose-response curves for each drug using primary BMDMs, primary murine embryonic fibroblasts (MEF), primary human endothelial cells (HUVECs), and the colon cancer cell line DLD-1 (Figure 3A). The inclusion of non-transformed primary cells enabled estimation of off-target toxicity in stromal and endothelial populations, while DLD-1 served as a benchmark for proliferative, transformed cells commonly used in drug screens. Due to their higher proliferative rate, DLD-1 cells were treated with higher drug concentrations than the other cell types. A safety margin was calculated by comparing the IC_90_ values of non-macrophage cell types to those of macrophages (Figure 3B, Table 1). Based on these data, we selected three compounds with the most favorable selectivity profiles for further characterization.

**Figure 3:**
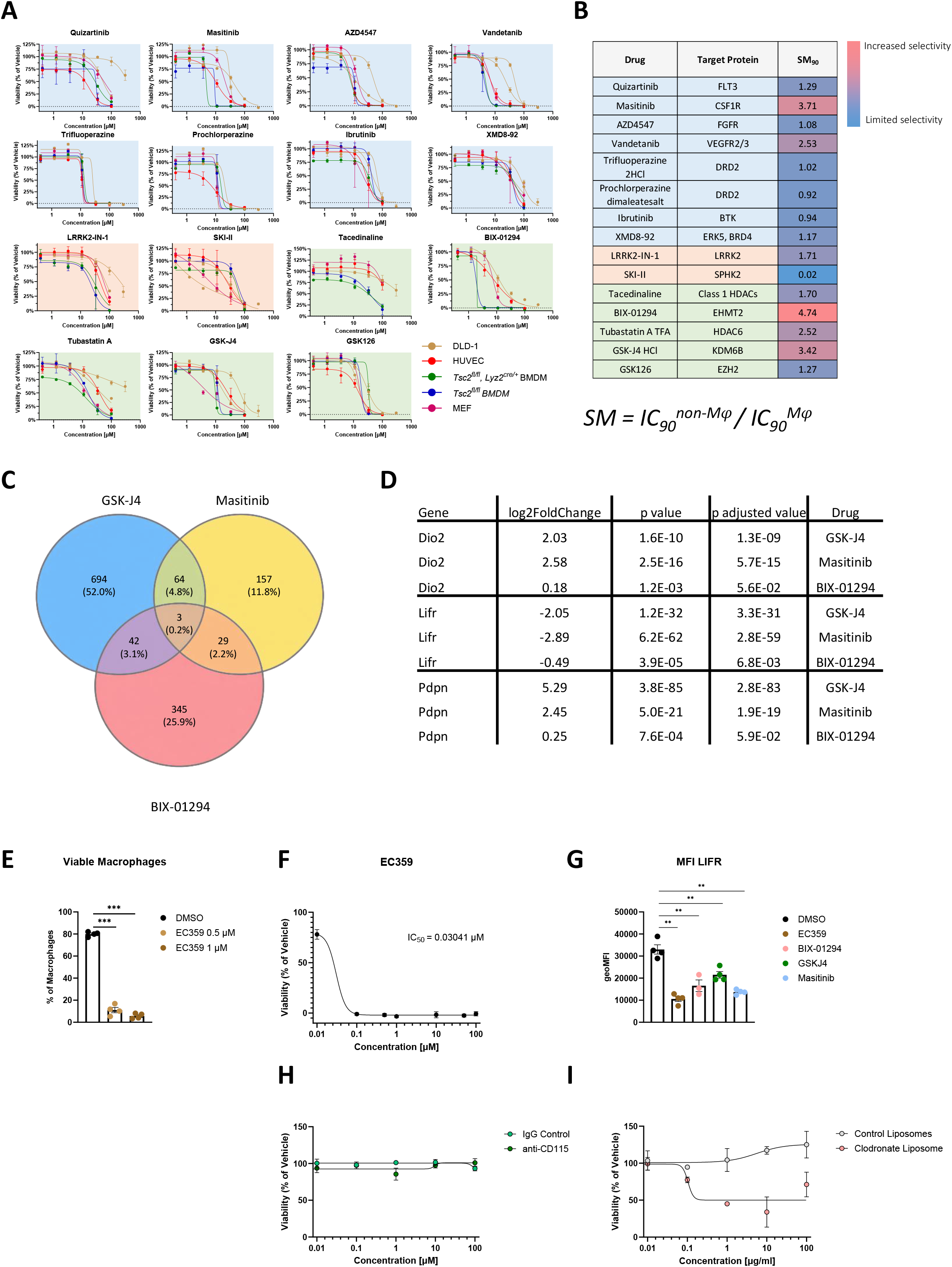
Three candidate drugs preferentially kill macrophages. **(A)** Dose–response curves for the indicated cell types treated with the specified compounds at concentrations ranging from 0.1 µM to 100 µM. **(B)** Table summarizing the identified compounds, their primary protein targets, and the calculated safety margins based on IC90 values. The formula used to calculate the safety margin is shown below the table. Red shading indicates higher macrophage selectivity, whereas blue shading indicates lower selectivity. Data represent mean ± SEM from three independent experiments with three technical replicates each. **(C)** Venn diagram showing differentially expressed genes (DEGs) following treatment with each compound and overlaps between treatments. Selection was based on adjusted p values (cutoff 0.1). **(D)** Table listing genes commonly regulated across all three compounds. **(E)** Percentage of viable macrophages following treatment with 0.5 or 1 µM of the LIFR antagonist EC359 for 16 hours. **(F)** Dose–response curve of bone marrow-derived macrophage viability following treatment with EC359 at concentrations ranging from 0.01 µM to 100 µM. **(G)** Mean fluorescence intensity (MFI) of LIFR on bone marrow-derived macrophages treated for 16 hours with EC359, BIX-01294, GSK-J4, or Masitinib. (H) Dose–response curve of bone marrow-derived macrophage viability following treatment with anti-CD115 antibody or IgG control ranging from 0.01 µM to 100 µM. (I) Dose–response curve of bone marrow-derived macrophage viability following treatment with clodronate liposomes or control liposomes ranging from 0.01 µg/ml to 100 µg/ml. Data represent mean ± SEM of four biological replicates. Statistical significance is indicated as ** p < 0.01, *** p < 0.001, **** p < 0.0001.

**Table 1:** IC90 concentrations and calculated safety margins for candidate drugs.

| Drug | Target Protein | Tsc2<sup>fl/fl</sup> BMDM | Tsc2<sup>fl/fl</sup>, Lyz2<sup>cre/+</sup> BMDM | DLD-1 | HUVEC | MEF | SM<sub>90</sub> |
| --- | --- | --- | --- | --- | --- | --- | --- |
| Quizartinib | FLT3 | 35.49 | 90.16 | 1.12E-230 | 45.71 | 135.2 | 1.29 |
| Masitinib | CSF1R | 9.7 | 4.744 | 236.5 | 35.99 | 39.01 | 3.71 |
| AZD4547 | FGFR | 13.24 | 14.39 | 108.7 | 14.27 | 68.58 | 1.08 |
| Vandetanib | VEGFR2/3 | 6.618 | 6.753 | 92.74 | 16.74 | 72.22 | 2.53 |
| Trifluoperazine 2HCl | DRD2 | 15.73 | 19.97 | 30.43 | 32.94 | 14.5 | 1.02 |
| Prochlorperazine dimaleatesalt | DRD2 | 14.82 | 15.68 | 31.35 | 18.16 | 15.17 | 0.92 |
| Ibrutinib | BTk | 56.75 | 67.48 | 95.38 | 53.34 | 110.6 | 0.94 |
| XMD8-92 | ERK5, BRD4 | 70.83 | 96.84 | 240.7 | 82.7 | 764.1 | 1.17 |
| LRRK2-IN-1 | LRRK2 | 67.77 | 77.29 | 7.197E-34 | 115.7 | 218 | 1.71 |
| SKI-II | SPHK2 | 115.6 | 83.82 | 133.7 | 110.8 | 2.491 | 0.02 |
| Tacedinaline | Class 1 HDACs | 180.7 | 307.9 | 4823 | N/A | 306.8 | 1.70 |
| BIX-01294 | EHMT2 | 2.605 | 3.36 | 77.68 | 54.71 | 12.34 | 4.74 |
| Tubastatin A TFA | HDAC6 | 50.85 | 80.36 | 138251 | 127.9 | 445.3 | 2.52 |
| GSK-J4 HCl | KDM6B | 15.99 | 11.48 | 156.9 | 117.5 | 54.65 | 3.42 |
| GSK126 | EZH2 | 23.44 | 40.1 | 73.09 | 46.57 | 29.79 | 1.27 |

**Table 2:**
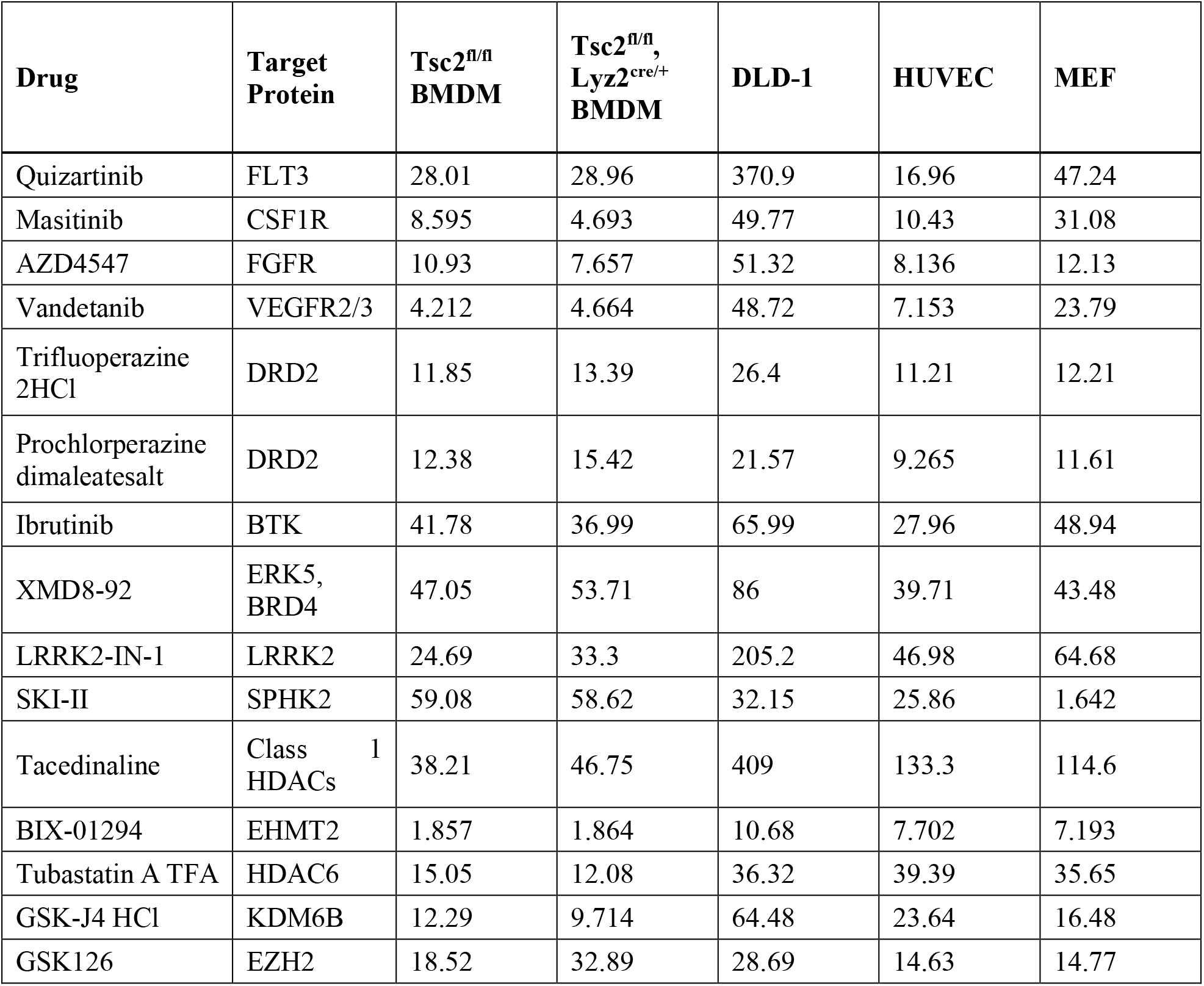
Derived IC50 concentrations for all drugs.

BIX-01294 inhibits the euchromatic histone lysine methyltransferase 2 (EHMT2)^36^, and EHMT2 loss promotes an M(LPS) phenotype in macrophages by impairing fatty acid uptake^37^. Masitinib is an inhibitor of CSF1R and c-KIT and has been used in preclinical studies to deplete microglia in amyotrophic lateral sclerosis models^38,39^. GSK-J4 inhibits jumonji domain-containing protein-3 (JMJD3, also known as KDM6B) and other KDM family members^40,41^ and has been previously linked to CSF1R signaling^42^.

To characterize the pathways targeted by these compounds, BMDMs were treated with each inhibitor and subjected to bulk RNA sequencing. GO pathway analysis revealed suppression of cell-cycle processes by BIX-01294 (Fig. S2A), and Masitinib (Figure S2C), whereas GSK-J4 primarily upregulated cell death-related pathways (Fig. S2B). Next, we compared the global gene expression profiles across all three treatments. Comparison of the global transcriptional profiles uncovered three genes consistently regulated across all treatments despite their distinct molecular targets (Figure 3C). Type II iodothyronine deiodinase (Dio2) and podoplanin (Pdpn) were upregulated, whereas leukemia inhibitory factor receptor (Lifr) was consistently downregulated (Figure 3D). While Dio2 and Pdpn are not known to regulate cell viability, Lifr has been implicated in apoptosis in several cancer cells, but its role in macrophages has not been defined^43–45^.

To evaluate whether LIFR contributes to macrophage survival, we treated BMDMs with the LIFR antagonist EC359^46^, which markedly reduced viability in a dose-dependent manner (Figure 3E, F). In agreement with the RNA sequencing data, all three compounds significantly decreased LIFR surface expression on BMDMs (Figure 3G). Although Lifr is expressed across multiple cell types, analysis of the Tabula muris dataset^47^, revealed that macrophages rank among the highest expressors, comprising two of the five top-expressing cell populations across tissues (Figure S3). Lastly, we used established methods for macrophage depletion both *in vitro* and *in vivo,* namely an anti-CD115 (CSF1R) antibody and clodronate liposomes, to compare their potency with that of the candidate compounds. While the anti-CSF1R antibody did not induce macrophage killing *in vitro* (Figure 3H), clodronate liposomes potently reduced macrophage viability (Figure 3I), although less strongly than the candidate compounds. Taken together, we identified three compounds with preferential cytotoxicity towards macrophages *in vitro* and show that all three converge on LIFR downregulation as a potential shared mechanism contributing to reduced macrophage viability.

### Large peritoneal macrophages are depleted by candidate drugs

To evaluate the *in vivo* effects of the selected compounds, we focused on macrophages in the peritoneal cavity under homeostatic conditions. This compartment contains two major subsets distinguished by size and surface marker expression: large peritoneal macrophages (LPMs) and small peritoneal macrophages (SPMs). SPMs are known precursors derived from monocytes that can differentiate into LPMs, following inflammatory or depletion-induced turnover. C57BL/6J mice received daily intraperitoneal injections of each compound for seven days, after which peritoneal exudate cells were analyzed by flow cytometry (Figure S4). All three treatments decreased LPM numbers, with significant reductions observed for BIX-01294 and Masitinib, whereas SPM abundance remained unchanged relative to vehicle controls (Figure 4A-C). Each treatment induced an increase in infiltrating monocytes (Figure 4D), consistent with compensatory recruitment following LPM depletion^48^.

**Figure 4:**
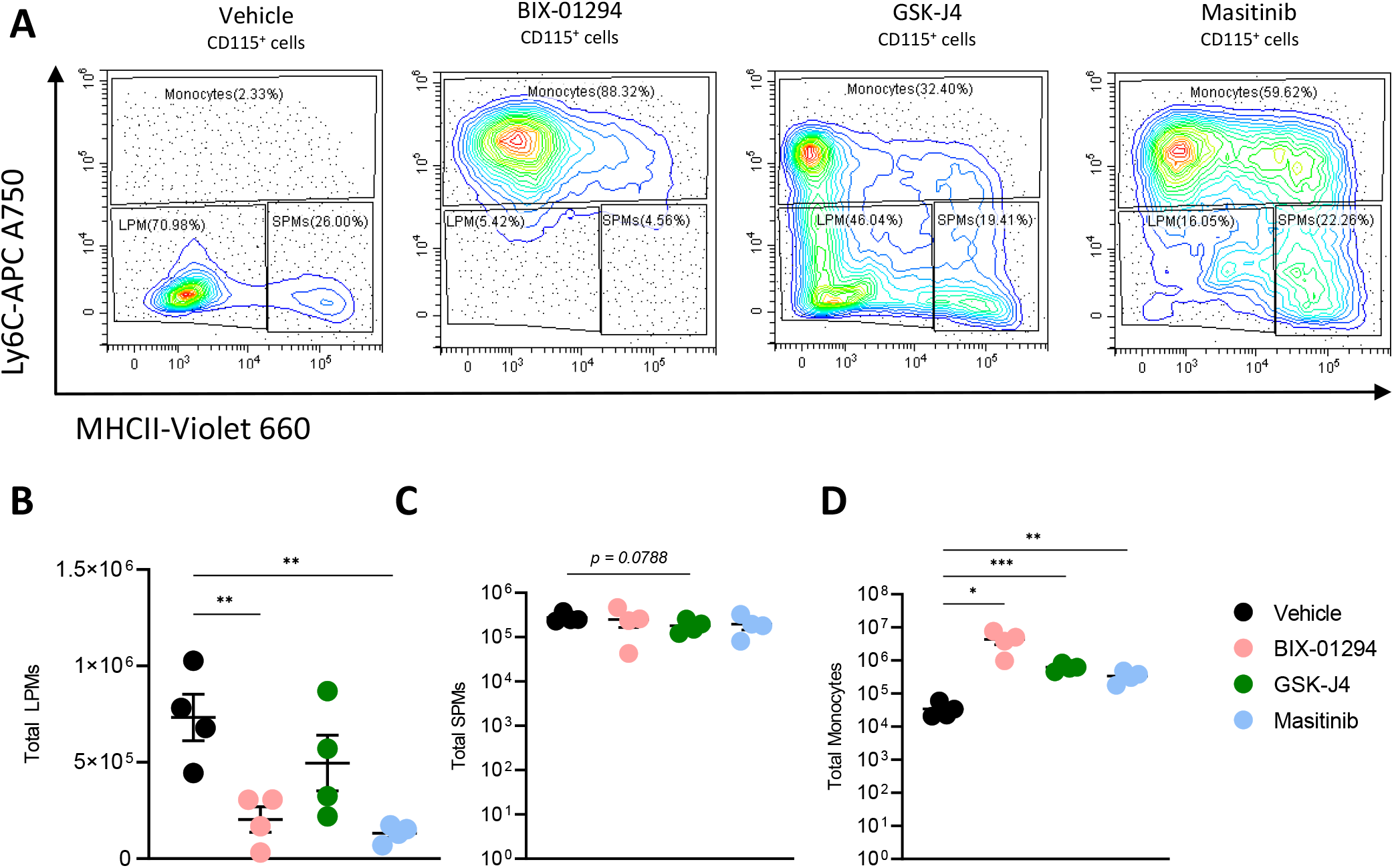
Candidate drugs deplete macrophages *in vivo*. **(A)** Representative flow cytometry contour plots showing peritoneal macrophage populations gated on 7-AAD^-^, CD45^+^, B220^-^, and CD115^+^ cells and distinguished based on Ly6C and MHC II expression. **(B–D)** Quantification of peritoneal macrophage populations shown in **(A)**. Cell numbers were normalized to the total number of cells recovered from the peritoneal cavity to calculate absolute numbers of each population. Data represent mean ± SEM of four biological replicates. Statistical significance is indicated as * p < 0.05, ** p < 0.01, *** p < 0.001.

We next assessed whether other immune cell types were affected. GSK-J4 reduced B-cell numbers, which is consistent with JMJD3 inhibition affecting BCL6, a key transcription factor required for B-cell maintenance (Figure S5A)^49,50^, and Masitinib caused a mild decrease as well (Figure S5A). Neutrophils were increased by BIX-01294 and slightly reduced by GSK-J4 (Figure S5B). Furthermore, to compare the candidate compounds with established macrophage-depletion approaches, we treated mice with clodronate liposomes or an anti-CD115 antibody and analyzed the peritoneal exudate. Both interventions led to a decrease in large peritoneal macrophages, while only the anti-CD115 antibody also led to a loss in small peritoneal macrophages (Figure S5C, D). Overall, these data show that the candidate drugs preferentially deplete LPMs *in vivo*, while sparing SPMs and most other immune populations, consistent with the preferential macrophage cytotoxicity observed *in vitro*.

### Macrophage activation syndrome disease progression is limited by candidate drugs

As a disease-relevant proof-of-concept, we employed a CpG-induced model of macrophage activation syndrome (MAS), a macrophage-driven pathology with minimal involvement of other immune cell types^22^. This preclinical model recapitulates characteristic features of human MAS, including hepatosplenomegaly and hepatic inflammation^8,22,51^. Mice received five injections of the TLR9 agonist CpG over ten days. Two days after the first injection, the candidate drugs were administered daily as a therapeutic intervention (Figure 5A). At the experimental endpoint, all three compounds significantly reduced splenomegaly (Figure 5B, C). Histological analysis of liver sections^52^ revealed reduced leukocyte infiltration, indicating attenuation of hepatic inflammation (Figure 5D). In addition, BIX-01294 and GSK-J4 reduced macrophage numbers in the liver (Figure 5E). To explore the translational relevance of these findings, we examined publicly available RNA sequencing datasets of monocytes isolated from patients suffering from sJIA, a condition that can progress to MAS^9^, as well as bulk RNA sequencing from whole blood of sJIA patients. In both datasets, KDM6A and KDM6B, the enzymatic targets of GSK-J4, were significantly upregulated compared healthy controls (Figure 5F). Together, these results indicate that the candidate drugs reduce key features of MAS pathology *in vivo*.

**Figure 5:**
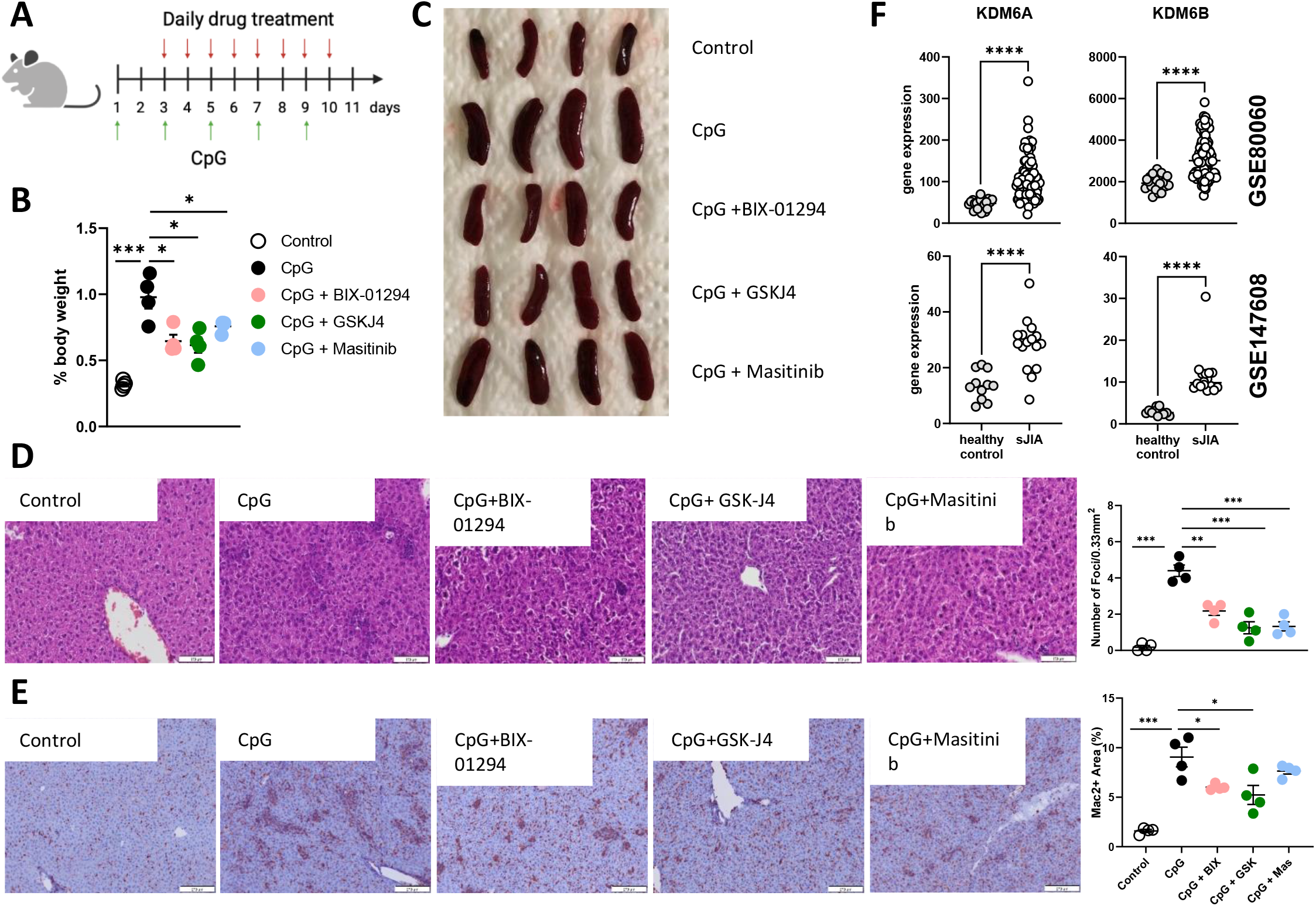
Macrophage activation syndrome severity is reduced by candidate drug treatment. **(A)** Experimental outline of the CpG-induced murine macrophage activation syndrome model. **(B)** Spleen weight expressed as a percentage of total body weight in control mice, CpG-injected mice, and CpG-injected mice treated with one of the candidate drugs (BIX-01294 [BIX], GSK-J4 [GSK], or Masitinib [Mas]). Data represent mean ± SEM of four biological replicates. **(C)** Representative images of spleens quantified in **(B)**. **(D)** Representative hematoxylin and eosin–stained liver sections from control mice, CpG-injected mice, and CpG-injected mice treated with candidate drugs, with quantification of leukocyte foci in the liver as described in the Methods. **(E)** Representative Mac-2 immunohistochemistry staining of liver sections from control mice, CpG-injected mice, and CpG-injected mice treated with candidate drugs, with quantification of Mac-2–positive area. Data represent mean ± SEM of four biological replicates. **(F)** Gene expression levels of the GSK-J4 target genes KDM6A and KDM6B in two publicly available RNA sequencing datasets (GSE80060 and GSE147608) from healthy controls and patients with systemic juvenile idiopathic arthritis (sJIA). GSE80060: healthy control n = 22, sJIA n = 104; GSE147608: healthy control n = 11, sJIA n = 16. Statistical significance is indicated as * p < 0.05, ** p < 0.01, **** p < 0.0001.

### Candidate drugs impair tumor growth and reduce macrophage numbers

To investigate the effects of the three compounds in a tumor setting, B16-F10 melanoma cells were injected subcutaneously into C57BL/6J mice, and treatment was initiated two days later to model a therapeutic intervention (Figure 6A). All three drugs significantly reduced tumor growth (Figure 6B - D). Immunofluorescence analysis of tumor sections revealed a marked decrease in F4/80-positive macrophages (Figure 6E, F). Most macrophages co-expressed CD206, consistent with a tumor-associated polarization profile (Figure 6G). The overall intensity of the maturation marker F4/80 was also reduced (Figure 6H).

**Figure 6:**
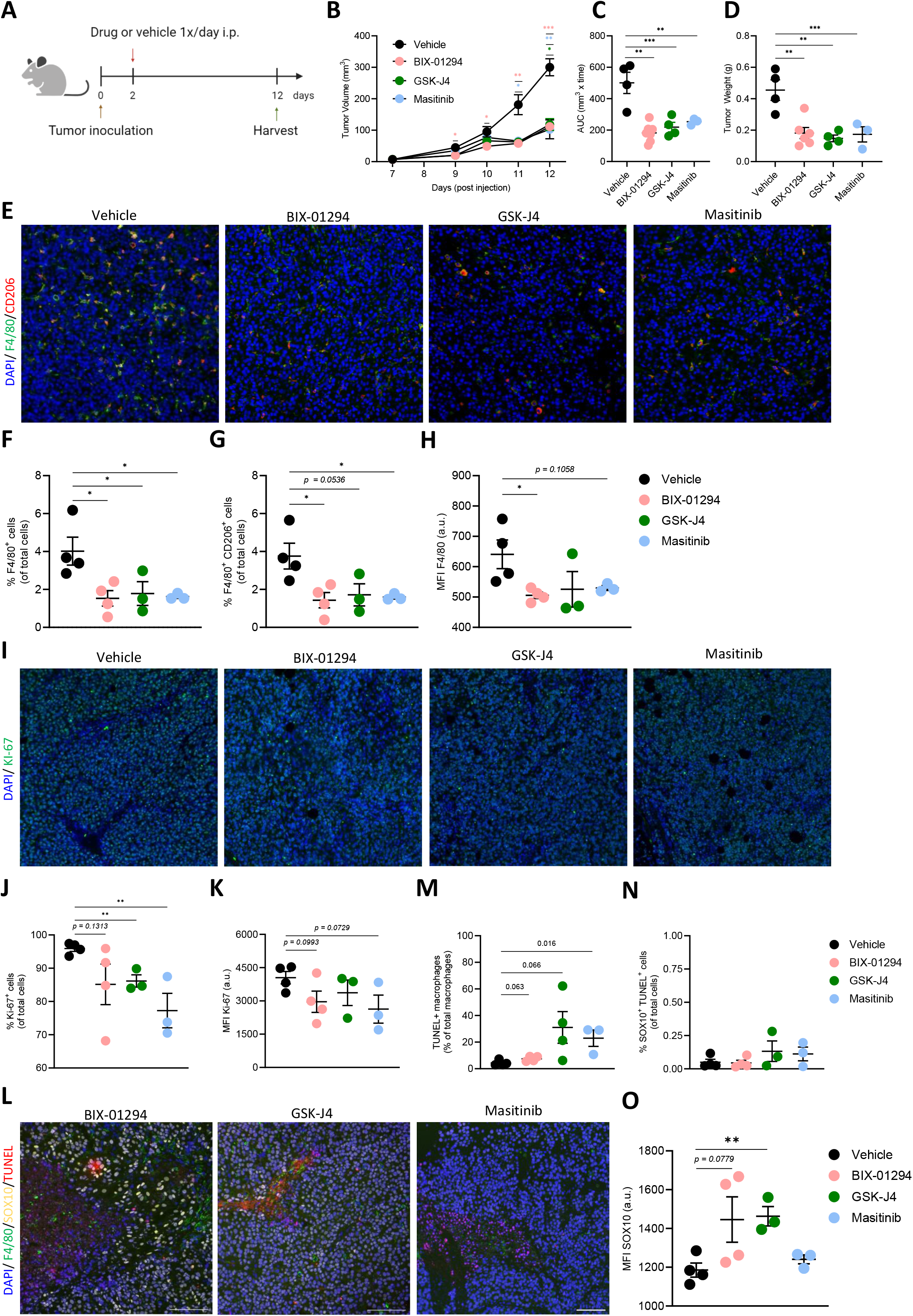
Candidate drugs target tumor-associated macrophages in a B16-F10 melanoma model. **(A)** Experimental outline of the B16-F10 melanoma tumor model and treatment regimen. **(B)** Tumor volume over time following tumor inoculation, quantified as described in the Methods. **(C)** Area under the curve (AUC) analysis of tumor volume for tumor-bearing mice. **(D)** Tumor weight measured at the experimental endpoint for all tumor-bearing mice. **(E)** Representative immunofluorescence images of tumor sections stained for DAPI, F4/80, and CD206. **(F)** Quantification of F4/80-positive cells as a percentage of total cells within the tumor. **(G)** Quantification of F4/80- and CD206-double-positive cells as a percentage of total cells within the tumor. Panels F and G were quantified from the same tumor sections, with the population quantified in G representing the CD206-positive subset of the F4/80-positive population quantified in F. **(H)** Mean fluorescence intensity (MFI) of F4/80 within the tumor. **(I)** Representative immunofluorescence images of tumor sections stained for DAPI and Ki-67. **(J)** Quantification of Ki-67–positive cells as a percentage of total cells within the tumor. **(K)** Mean fluorescence intensity (MFI) of Ki-67 within the tumor. **(L)** Representative immunofluorescence images of tumor sections stained for DAPI, F4/80, SOX10, and TUNEL. **(M)** Quantification of TUNEL-positive macrophages as a percentage of total macrophages within the tumor. **(N)** Quantification of TUNEL- and SOX10-double-positive cells as a percentage of total cells within the tumor. **(O)** Mean fluorescence intensity (MFI) of SOX10 within the tumor. All quantifications represent mean ± SEM of three to seven biological replicates. Statistical significance is indicated as * p < 0.05, ** p < 0.01, *** p < 0.001.

Staining tumor sections for the proliferation marker Ki-67 showed a significant reduction in total proliferating cells within the tumor microenvironment (Figure 6I - K). Apoptosis was assessed by staining for chromatin fragmentation by fluorescently labelling DNA nicks, and all three compounds showed an increase in TUNEL+ apoptotic macrophages (Figure 6L, M). In contrast, SOX10-postive melanoma cells did not show increased apoptosis in tumors of the treated animals (Figure 6N). Notably, SOX10 expression was increased in groups treated with BIX-01294 and GSK-J4, in line with previous reports linking SOX10 upregulation to altered melanoma cell states^53^ (Figure 6O). Neither endothelial nor CD8+ T-cell numbers were significantly altered (Figure S6), consistent with a contribution of macrophage depletion to the observed anti-tumor effect (Figure S6A, B).

To extend these findings, we analyzed publicly available bulk RNA sequencing data from melanoma patients. Expression of the target genes of Masitinib, BIX-01294, and GSK-J4 positively correlated with the macrophage-associated genes SIGLEC1 and MRC1 (Figure S7A, B). Together, these results indicate that the compounds reduce tumor growth and preferentially induce macrophage apoptosis within the tumor microenvironment.

Finally, we aimed to evaluate the therapeutic potential of one compound in a genetically engineered *K-ras* ^LSLG12D^:*p53* ^fl/fl^autochthonous lung cancer model. In this model, intratracheal administration of a Cre-expressing adenovirus activates oncogenic K-Ras and deletes p53 in type II alveolar cells, initiating tumor formation. Given its regulatory approval history, Masitinib was selected for evaluation in this model. One day after adenoviral instillation, mice were treated daily with Masitinib or vehicle control for 82 days (Figure 7A). Masitinib significantly reduced overall tumor burden (Figure 7B – D) and decreased individual nodule size (Figure 7E). There was also a trend toward reduced macrophage numbers in the tumors of the Masitinib-treated mice (Figure 7F – H). Collectively, these experiments demonstrate the *in vivo* efficacy of the identified compounds in both inflammatory and oncological disease contexts and support their potential as macrophage-targeted therapeutic agents.

**Figure 7:**
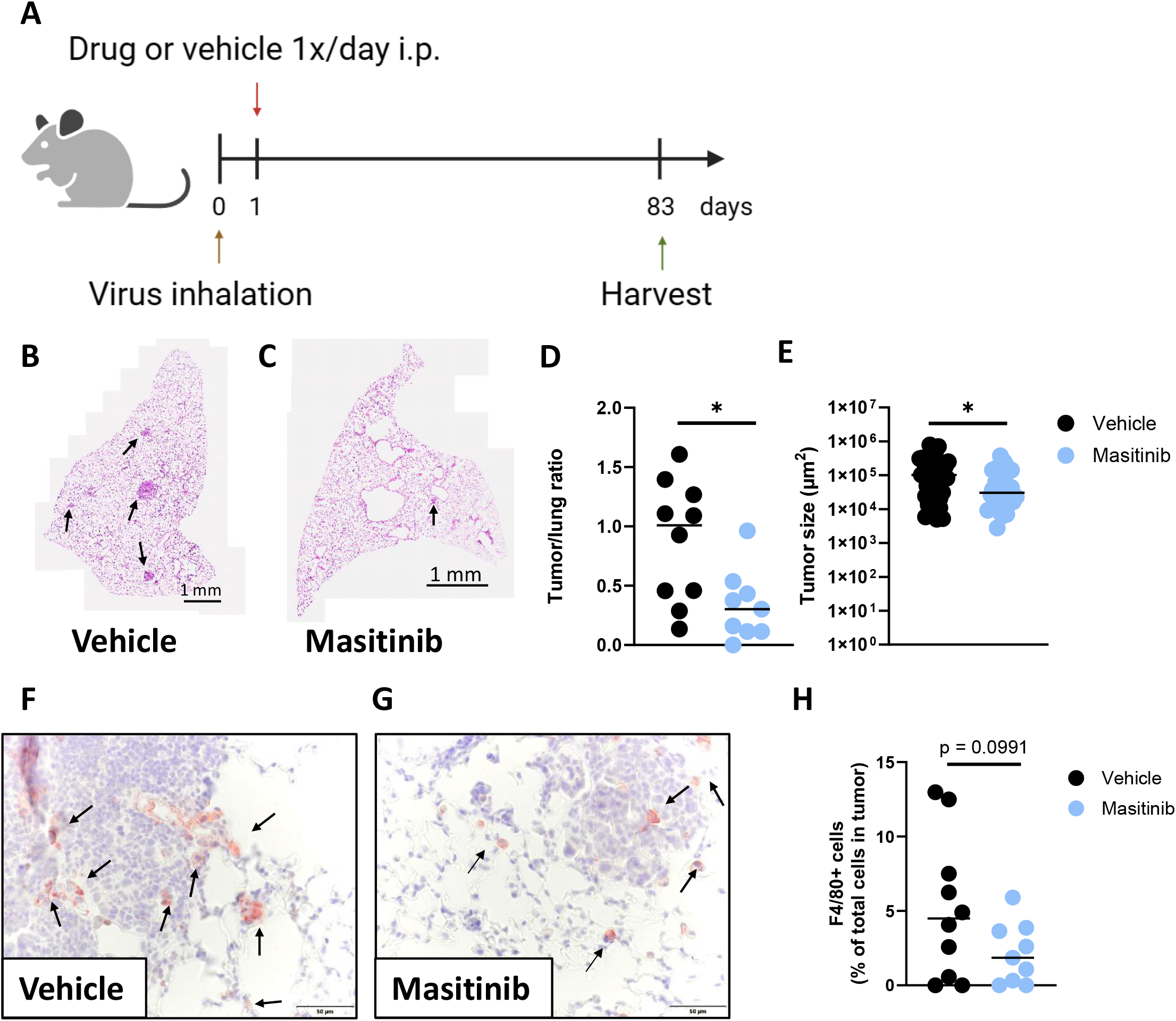
Masitinib dampens tumor growth in an autochthonous lung cancer model. **(A)** Experimental outline of the K-ras-driven autochthonous lung cancer model and Masitinib treatment regimen. **(B, C)** Representative hematoxylin and eosin–stained lung sections from vehicle-treated and Masitinib-treated mice. Arrows indicate tumor nodules. **(D)** Quantification of tumor burden expressed as the ratio of tumor area to total lung area (vehicle control n = 10; Masitinib-treated n = 9). **(E)** Quantification of individual tumor nodule size (vehicle control n = 39 tumor nodules; Masitinib-treated n = 33 tumor nodules). **(F, G)** Representative images of F4/80 immunohistochemistry staining of lung sections from vehicle-treated and Masitinib-treated mice. Arrows indicate F4/80-positive macrophages. **(H)** Quantification of F4/80-positive macrophages per tumor as percentage of total cells per tumor. For all quantifications, horizontal lines indicate the mean. Statistical significance is indicated as * p < 0.05.

## Discussion

The present study identifies pharmacological vulnerabilities in macrophage survival using a high-throughput cell death screening approach combined with *in vitro* and *in vivo* validation. Although macrophages are central contributors to inflammatory and malignant diseases, therapeutic strategies that directly reduce macrophage numbers remain limited. Blockade of CSF1R signaling has demonstrated that macrophage depletion can be clinically meaningful^1,54^, yet its efficacy is frequently constrained by compensatory pathways and incomplete responses. Here, we identify three structurally distinct small molecules that consistently reduce macrophage viability *in vitro* and macrophage abundance in disease-relevant *in vivo* settings. Despite their distinct primary targets, all three compounds were associated with reduced LIFR expression, suggesting convergence on shared macrophage survival pathways. *In vivo*, treatment with these agents ameliorated macrophage activation syndrome (MAS) pathology and limited tumor growth, supporting their therapeutic potential. Together, these findings demonstrate the utility of this screening approach for identifying macrophage-depleting compounds and highlight LIFR-associated signaling as a regulator of macrophage survival regulation.

The three compounds identified act through distinct, previously described molecular targets. BIX-01294 inhibits the histone methyltransferase EHMT2, which has been implicated in metabolic regulation in macrophages, including control of fatty acid uptake through CD36^37^. GSK-J4 inhibits the histone demethylases KDM6B and KDM6A and has been linked to macrophage activation-associated transcriptional programs^42^. Masitinib inhibits CSF1R and c-KIT and has been evaluated in preclinical and clinical studies of amyotrophic lateral sclerosis^39,55^. Despite these distinct targets, all three compounds reduced macrophage viability, predominantly through non-lytic cell death pathways *in vitro*, with GSK-J4 additionally showing features compatible with lytic activity.

Beyond their known targets, leukemia inhibitory factor receptor (LIFR) emerged as a shared pathway modulated by all three compounds. LIFR has been studied as a therapeutic target in cancer due to its role in promoting tumor cell proliferation, migration, and survival via autocrine LIF signaling ^56^. More recently, LIFR signaling has been implicated in promoting immunosuppressive and tumor-supportive macrophage functions^57,58^. In line with these observations, treatment with all three compounds reduced the abundance of F4/80- and CD206-positive macrophages in tumor tissue. Although none of the compounds had previously been linked directly to LIFR regulation, epigenetic mechanisms have been shown to influence LIFR expression, providing a plausible connection to the epigenetic modulators BIX-01294 and GSK-J4^59,60^. In addition, LIFR signaling enhances CSF1 consumption through CSF1R-dependent mechanisms^57^, offering a potential link between LIFR modulation and the activity of Masitinib.

*In vitro* experiments using both wild-type and Tsc2-deficient bone marrow-derived macrophages provided further insight into compound sensitivity under conditions of altered signaling activity. Tsc2-deficient macrophages display a hyperactivated phenotype, and while all three compounds reduced viability of both macrophage populations after prolonged exposure, Tsc2-deficient macrophages were affected at earlier time points. This suggests that macrophages with heightened activation states may be particularly sensitive to disruption of the survival pathways targeted by these compounds. Analysis of the Tabula Muris dataset^47^ showed that several established targets of the candidate compounds, particularly CSF1R and KDM6B, are enriched in myeloid populations across tissues (data not shown), suggesting that differential target expression may contribute to preferential macrophage sensitivity.

The CpG-induced MAS model served as a disease-relevant *in vivo* proof of concept. In this setting, all three compounds ameliorated key pathological features, including splenomegaly and hepatic inflammation. Current therapeutic approaches for MAS largely focus on blocking individual cytokines that drive disease pathology^9,51^. However, a subset of patients remains refractory to cytokine-targeted therapies^10^. Given the broad and dysregulated activation state of macrophages in MAS^7,8^, directly reducing pathogenic macrophage populations may represent a useful alternative to cytokine-targeted approaches.

In cancer models, the candidate compounds significantly impaired tumor growth in a transplanted melanoma model and Masitinib showed efficacy in an autochthonous lung cancer model. In the B16 melanoma model, treatment reduced macrophage abundance within tumors and increased apoptotic macrophage death, whereas tumor cells themselves did not display enhanced apoptosis. This suggests that reduced tumor growth is driven, at least in part, by altered macrophage-tumor interactions rather than direct toxicity to cancer cells. These findings are consistent with the well-established role of macrophages in promoting tumor growth, immune suppression, and therapy resistance within the tumor microenvironment.

Targeting macrophages has long been explored as a therapeutic strategy in cancer, most prominently through inhibition of CSF1R signaling^54,61^. While CSF1R blockade has shown clinical efficacy in selected tumor entities such as tenosynovial giant cell tumor^62^, responses in many solid tumors have been limited or variable^54^. One challenge is the capacity of macrophages to engage alternative survival and proliferative pathways, thereby limiting the durability of CSF1R-directed therapies^63^. The compounds identified here reduce macrophage numbers and tumor growth through mechanisms distinct from CSF1R blockade alone, supporting the concept that targeting macrophage survival pathways may overcome some of these limitations.

The compounds identified here reduced macrophage numbers without significantly altering endothelial cells or CD8-positive T-cell abundance within tumors, indicating a relatively focused effect on macrophage populations. As macrophages are key contributors to immunosuppression in the tumor microenvironment, macrophage-depleting strategies have been proposed as combination partners for immune checkpoint blockade^64,65^. The lack of overt effects on adaptive immune cells suggests that the compounds described here could complement checkpoint inhibitors through non-overlapping mechanisms, a concept that may be particularly relevant for patients who do not respond to checkpoint blockade alone^66^.

The efficacy of Masitinib in an autochthonous K-ras–driven lung cancer model further supports the relevance of macrophage-directed strategies across tumor types. Lung cancer patients increasingly receive immune checkpoint inhibitors as standard-of-care therapy^67^, yet a substantial proportion fails to respond. Macrophage-associated gene expression signatures have been shown to predict response to immunotherapy^68^, highlighting macrophages as key modulators of therapeutic outcome. In this context, macrophage-depleting approaches may represent a rational combination strategy with immune checkpoint blockade, particularly in tumors characterized by macrophage-driven immune suppression^64,65,69^.

In summary, we present a screening-based pipeline that identified three small molecules capable of preferentially reducing macrophage viability *in vitro* and macrophage abundance *in vivo*. These compounds were associated with downregulation of LIFR, whose inhibition impaired macrophage survival. Together, our findings support macrophage depletion as a viable therapeutic strategy in inflammatory and oncological disease settings and provide a framework for the identification of additional macrophage-targeted agents.

## Acknowledgments

We would like to thank Jaqueline Horvath for help with mouse experiments and Michael Machtinger for histology work. Next-generation sequencing (and initial data analysis) was performed by the Biomedical Sequencing Facility at CeMM Research Center for Molecular Medicine of the Austrian Academy of Sciences.

## Author contributions

Conceptualization: SF, PE, TW

Methodology: SF, PE, MT, M. Mazic, HKM, AV, LD, RS, VSA, AG, CT, AK, SK

Investigation: SF, PE, MT, M. Mazic, HKM, AV, LD, RS, VSA, AG, CT, AK, SK

Visualization: SF, PE, TW

Funding acquisition: HPM, EC, HD, MH, TR, SF, TW

Supervision: TW

Writing – original draft: SF, PE, TW

Writing – review & editing: all authors

## References

1 Park MD, Silvin A, Ginhoux F, Merad M. Macrophages in health and disease. Cell 2022; 185: 4259–4279.

2 Fritsch SD, Sukhbaatar N, Gonzales K, Sahu A, Tran L, Vogel A et al. Metabolic support by macrophages sustains colonic epithelial homeostasis. Cell Metab 2023; 35: 1931–1943.e8.

3 Ginhoux F, Schultze JL, Murray PJ, Ochando J, Biswas SK. New insights into the multidimensional concept of macrophage ontogeny, activation and function. Nat Immunol 2016; 17: 34–40.

4 Mass E, Nimmerjahn F, Kierdorf K, Schlitzer A. Tissue-specific macrophages: how they develop and choreograph tissue biology. Nat Rev Immunol 2023; 23: 563–579.

5 Honold L, Nahrendorf M. Resident and Monocyte-Derived Macrophages in Cardiovascular Disease. Circ Res 2018; 122: 113–127.

6 Saha P, Ettel P, Weichhart T. Leveraging macrophage metabolism for anticancer therapy: opportunities and pitfalls. Trends Pharmacol Sci 2024; 45: 335–349.

7 Grom AA, Horne A, De Benedetti F. Macrophage activation syndrome in the era of biologic therapy. Nat Rev Rheumatol 2016; 12: 259–268.

8 Crayne CB, Albeituni S, Nichols KE, Cron RQ. The immunology of macrophage activation syndrome. Front Immunol 2019; 10: 119.

9 McGonagle D, Ramanan AV, Bridgewood C. Immune cartography of macrophage activation syndrome in the COVID-19 era. Nat Rev Rheumatol 2021; 17: 145–157.

10 Grom AA, Ilowite NT, Pascual V, Brunner HI, Martini A, Lovell D et al. Rate and clinical presentation of macrophage activation syndrome in patients with systemic juvenile idiopathic arthritis treated with canakinumab. Arthritis Rheumatol 2016; 68: 218–228.

11 Nigrovic PA, Mannion M, Prince FHM, Zeft A, Rabinovich CE, van Rossum MAJ et al. Anakinra as first-line disease-modifying therapy in systemic juvenile idiopathic arthritis: report of forty-six patients from an international multicenter series. Arthritis Rheum 2011; 63: 545–555.

12 Li M, He L, Zhu J, Zhang P, Liang S. Targeting tumor-associated macrophages for cancer treatment. Cell Biosci 2022; 12: 85.

13 Awad RM, De Vlaeminck Y, Maebe J, Goyvaerts C, Breckpot K. Turn back the time: targeting tumor infiltrating myeloid cells to revert cancer progression. Front Immunol 2018; 9: 1977.

14 Haas L, Obenauf AC. Allies or Enemies-The Multifaceted Role of Myeloid Cells in the Tumor Microenvironment. Front Immunol 2019; 10: 2746.

15 Sabharwal SS, Schumacker PT. Mitochondrial ROS in cancer: initiators, amplifiers or an Achilles’ heel? Nat Rev Cancer 2014; 14: 709–721.

16 Quail DF, Joyce JA. Molecular Pathways: Deciphering Mechanisms of Resistance to Macrophage-Targeted Therapies. Clin Cancer Res 2017; 23: 876–884.

17 Ostuni R, Kratochvill F, Murray PJ, Natoli G. Macrophages and cancer: from mechanisms to therapeutic implications. Trends Immunol 2015; 36: 229–239.

18 Ngambenjawong C, Gustafson HH, Pun SH. Progress in tumor-associated macrophage (TAM)-targeted therapeutics. Adv Drug Deliv Rev 2017; 114: 206–221.

19 Ponzoni M, Pastorino F, Di Paolo D, Perri P, Brignole C. Targeting macrophages as a potential therapeutic intervention: impact on inflammatory diseases and cancer. Int J Mol Sci 2018; 19. doi:10.3390/ijms19071953.

20 Goswami S, Raychaudhuri D, Singh P, Natarajan SM, Chen Y, Poon C et al. Myeloid-specific KDM6B inhibition sensitizes glioblastoma to PD1 blockade. Nat Cancer 2023; 4: 1455–1473.

21 Linke M, Pham HTT, Katholnig K, Schnöller T, Miller A, Demel F et al. Chronic signaling via the metabolic checkpoint kinase mTORC1 induces macrophage granuloma formation and marks sarcoidosis progression. Nat Immunol 2017; 18: 293–302.

22 Behrens EM, Canna SW, Slade K, Rao S, Kreiger PA, Paessler M et al. Repeated TLR9 stimulation results in macrophage activation syndrome-like disease in mice. J Clin Invest 2011; 121: 2264–2277.

23 Breitenecker K, Homolya M, Luca AC, Lang V, Trenk C, Petroczi G et al. Down-regulation of A20 promotes immune escape of lung adenocarcinomas. Sci Transl Med 2021; 13. doi:10.1126/scitranslmed.abc3911.

24 Moll HP, Pranz K, Musteanu M, Grabner B, Hruschka N, Mohrherr J et al. Afatinib restrains K-RAS-driven lung tumorigenesis. Sci Transl Med 2018; 10. doi:10.1126/scitranslmed.aao2301.

25 Jackson EL, Olive KP, Tuveson DA, Bronson R, Crowley D, Brown M et al. The differential effects of mutant p53 alleles on advanced murine lung cancer. Cancer Res 2005; 65: 10280–10288.

26 Weichhart T, Costantino G, Poglitsch M, Rosner M, Zeyda M, Stuhlmeier KM et al. The TSC-mTOR signaling pathway regulates the innate inflammatory response. Immunity 2008; 29: 565–577.

27 Byles V, Covarrubias AJ, Ben-Sahra I, Lamming DW, Sabatini DM, Manning BD et al. The TSC-mTOR pathway regulates macrophage polarization. Nat Commun 2013; 4: 2834.

28 Huang SC-C, Smith AM, Everts B, Colonna M, Pearce EL, Schilling JD et al. Metabolic Reprogramming Mediated by the mTORC2-IRF4 Signaling Axis Is Essential for Macrophage Alternative Activation. Immunity 2016; 45: 817–830.

29 Licciardello MP, Ringler A, Markt P, Klepsch F, Lardeau C-H, Sdelci S et al. A combinatorial screen of the CLOUD uncovers a synergy targeting the androgen receptor. Nat Chem Biol 2017; 13: 771–778.

30 Xie Z, Bailey A, Kuleshov MV, Clarke DJB, Evangelista JE, Jenkins SL et al. Gene Set Knowledge Discovery with Enrichr. Curr Protoc 2021; 1: e90.

31 Kropiwnicki E, Evangelista JE, Stein DJ, Clarke DJB, Lachmann A, Kuleshov MV et al. Drugmonizome and Drugmonizome-ML: integration and abstraction of small molecule attributes for drug enrichment analysis and machine learning. Database (Oxford) 2021; 2021. doi:10.1093/database/baab017.

32 Magtanong L, Ko PJ, Dixon SJ. Emerging roles for lipids in non-apoptotic cell death. Cell Death Differ 2016; 23: 1099–1109.

33 Smirnov P, Kofia V, Maru A, Freeman M, Ho C, El-Hachem N et al. PharmacoDB: an integrative database for mining in vitro anticancer drug screening studies. Nucleic Acids Res 2018; 46: D994–D1002.

34 Yang J, Liu Z, Wang C, Yang R, Rathkey JK, Pinkard OW et al. Mechanism of gasdermin D recognition by inflammatory caspases and their inhibition by a gasdermin D-derived peptide inhibitor. Proc Natl Acad Sci USA 2018; 115: 6792–6797.

35 Vandenabeele P, Galluzzi L, Vanden Berghe T, Kroemer G. Molecular mechanisms of necroptosis: an ordered cellular explosion. Nat Rev Mol Cell Biol 2010; 11: 700–714.

36 Kubicek S, O’Sullivan RJ, August EM, Hickey ER, Zhang Q, Teodoro ML et al. Reversal of H3K9me2 by a small-molecule inhibitor for the G9a histone methyltransferase. Mol Cell 2007; 25: 473–481.

37 Wang X, Chen S, He J, Chen W, Ding Y, Huang J et al. Histone methyltransferases G9a mediated lipid-induced M1 macrophage polarization through negatively regulating CD36. Metab Clin Exp 2021; 114: 154404.

38 Trias E, Ibarburu S, Barreto-Núñez R, Babdor J, Maciel TT, Guillo M et al. Post-paralysis tyrosine kinase inhibition with masitinib abrogates neuroinflammation and slows disease progression in inherited amyotrophic lateral sclerosis. J Neuroinflammation 2016; 13: 177.

39 Harrison JM, Rafuse VF. Muscle fiber-type specific terminal Schwann cell pathology leads to sprouting deficits following partial denervation in SOD1G93A mice. Neurobiol Dis 2020; 145: 105052.

40 Heinemann B, Nielsen JM, Hudlebusch HR, Lees MJ, Larsen DV, Boesen T et al. Inhibition of demethylases by GSK-J1/J4. Nature 2014; 514: E1–2.

41 Kruidenier L, Chung C, Cheng Z, Liddle J, Che K, Joberty G et al. A selective jumonji H3K27 demethylase inhibitor modulates the proinflammatory macrophage response. Nature 2012; 488: 404–408.

42 Satoh T, Takeuchi O, Vandenbon A, Yasuda K, Tanaka Y, Kumagai Y et al. The Jmjd3-Irf4 axis regulates M2 macrophage polarization and host responses against helminth infection. Nat Immunol 2010; 11: 936–944.

43 Tang W, Ramasamy K, Pillai SMA, Santhamma B, Konda S, Pitta Venkata P et al. LIF/LIFR oncogenic signaling is a novel therapeutic target in endometrial cancer. Cell Death Discov 2021; 7: 216.

44 Yao F, Deng Y, Zhao Y, Mei Y, Zhang Y, Liu X et al. A targetable LIFR-NF-κB-LCN2 axis controls liver tumorigenesis and vulnerability to ferroptosis. Nat Commun 2021; 12: 7333.

45 Li M, Viswanadhapalli S, Santhamma B, Pratap UP, Luo Y, Liu J, et al. LIFR inhibition enhances the therapeutic efficacy of HDAC inhibitors in triple negative breast cancer. Commun Biol 2021; 4: 1235.

46 Viswanadhapalli S, Luo Y, Sareddy GR, Santhamma B, Zhou M, Li M et al. EC359: A First-in-Class Small-Molecule Inhibitor for Targeting Oncogenic LIFR Signaling in Triple-Negative Breast Cancer. Mol Cancer Ther 2019; 18: 1341–1354.

47 Tabula Muris Consortium, Overall coordination, Logistical coordination, Organ collection and processing, Library preparation and sequencing, Computational data analysis et al. Single-cell transcriptomics of 20 mouse organs creates a Tabula Muris. Nature 2018; 562: 367–372.

48 Han J, Gallerand A, Erlich EC, Helmink BA, Mair I, Li X et al. Human serous cavity macrophages and dendritic cells possess counterparts in the mouse with a distinct distribution between species. Nat Immunol 2024; 25: 155–165.

49 Zhang Y, Shen L, Stupack DG, Bai N, Xun J, Ren G et al. JMJD3 promotes survival of diffuse large B-cell lymphoma subtypes via distinct mechanisms. Oncotarget 2016; 7: 29387–29399.

50 Huppertz S, Senger K, Brown A, Leins H, Eiwen K, Mulaw MA et al. KDM6A, a histone demethylase, regulates stress hematopoiesis and early B-cell differentiation. Exp Hematol 2021; 99: 32–43.e13.

51 Ravelli A, Minoia F, Davì S, Horne A, Bovis F, Pistorio A et al. 2016 classification criteria for macrophage activation syndrome complicating systemic juvenile idiopathic arthritis: A european league against rheumatism/american college of rheumatology/paediatric rheumatology international trials organisation collaborative initiative. Ann Rheum Dis 2016; 75: 481–489.

52 Liang W, Menke AL, Driessen A, Koek GH, Lindeman JH, Stoop R et al. Establishment of a general NAFLD scoring system for rodent models and comparison to human liver pathology. PLoS ONE 2014; 9: e115922.

53 Capparelli C, Purwin TJ, Glasheen M, Caksa S, Tiago M, Wilski N et al. Targeting SOX10-deficient cells to reduce the dormant-invasive phenotype state in melanoma. Nat Commun 2022; 13: 1381.

54 Mantovani A, Allavena P, Marchesi F, Garlanda C. Macrophages as tools and targets in cancer therapy. Nat Rev Drug Discov 2022; 21: 799–820.

55 Trias E, Kovacs M, King PH, Si Y, Kwon Y, Varela V et al. Schwann cells orchestrate peripheral nerve inflammation through the expression of CSF1, IL-34, and SCF in amyotrophic lateral sclerosis. Glia 2020; 68: 1165–1181.

56 Viswanadhapalli S, Dileep KV, Zhang KYJ, Nair HB, Vadlamudi RK. Targeting LIF/LIFR signaling in cancer. Genes Dis 2022; 9: 973–980.

57 Duluc D, Delneste Y, Tan F, Moles M-P, Grimaud L, Lenoir J et al. Tumor-associated leukemia inhibitory factor and IL-6 skew monocyte differentiation into tumor-associated macrophage-like cells. Blood 2007; 110: 4319–4330.

58 Pascual-García M, Bonfill-Teixidor E, Planas-Rigol E, Rubio-Perez C, Iurlaro R, Arias A et al. LIF regulates CXCL9 in tumor-associated macrophages and prevents CD8+ T cell tumor-infiltration impairing anti-PD1 therapy. Nat Commun 2019; 10: 2416.

59 Lv S, Ji L, Chen B, Liu S, Lei C, Liu X et al. Histone methyltransferase KMT2D sustains prostate carcinogenesis and metastasis via epigenetically activating LIFR and KLF4. Oncogene 2018; 37: 1354–1368.

60 Lu B, He Y, He J, Wang L, Liu Z, Yang J et al. Epigenetic Profiling Identifies LIF as a Super-enhancer-Controlled Regulator of Stem Cell-like Properties in Osteosarcoma. Mol Cancer Res 2020; 18: 57–67.

61 Chen Q, Zhang L, Li L, Tan M, Liu W, Liu S et al. Cancer cell membrane-coated nanoparticles for bimodal imaging-guided photothermal therapy and docetaxel-enhanced immunotherapy against cancer. J Nanobiotechnology 2021; 19: 449.

62 Tap WD, Gelderblom H, Palmerini E, Desai J, Bauer S, Blay J-Y et al. Pexidartinib versus placebo for advanced tenosynovial giant cell tumour (ENLIVEN): a randomised phase 3 trial. Lancet 2019; 394: 478–487.

63 Jenkins SJ, Ruckerl D, Thomas GD, Hewitson JP, Duncan S, Brombacher F et al. IL-4 directly signals tissue-resident macrophages to proliferate beyond homeostatic levels controlled by CSF-1. J Exp Med 2013; 210: 2477–2491.

64 Zhu Y, Knolhoff BL, Meyer MA, Nywening TM, West BL, Luo J et al. CSF1/CSF1R blockade reprograms tumor-infiltrating macrophages and improves response to T-cell checkpoint immunotherapy in pancreatic cancer models. Cancer Res 2014; 74: 5057–5069.

65 Mao Y, Eissler N, Blanc KL, Johnsen JI, Kogner P, Kiessling R. Targeting suppressive myeloid cells potentiates checkpoint inhibitors to control spontaneous neuroblastoma. Clin Cancer Res 2016; 22: 3849–3859.

66 Haddad AF, Young JS, Gill S, Aghi MK. Resistance to immune checkpoint blockade: Mechanisms, counter-acting approaches, and future directions. Semin Cancer Biol 2022; 86: 532–541.

67 Gandhi L, Rodríguez-Abreu D, Gadgeel S, Esteban E, Felip E, De Angelis F et al. Pembrolizumab plus Chemotherapy in Metastatic Non-Small-Cell Lung Cancer. N Engl J Med 2018; 378: 2078–2092.

68 Xiong D, Wang Y, You M. A gene expression signature of TREM2hi macrophages and γδ T cells predicts immunotherapy response. Nat Commun 2020; 11: 5084.

69 Cannarile MA, Weisser M, Jacob W, Jegg A-M, Ries CH, Rüttinger D. Colony-stimulating factor 1 receptor (CSF1R) inhibitors in cancer therapy. J Immunother Cancer 2017; 5: 53.

