## Supplementary Material for "A cell death screen identifies macrophage-depleting agents with therapeutic potential"

1 **Supplementary Data for:**
2 **A cell death screen identifies macrophage-depleting agents with**
3 **therapeutic potential**
4 **Stephan Forisch and Paul Ettel et al.**

5

6 File contains:

7 *Materials*

8 *Figs. S1 to S7*

9

10 **Materials**

| **Reagent or Resource** | **Source** | **Catalogue Number** |
| --- | --- | --- |
| **Antibodies** |  |  |
| Rabbit anti-GSDMD | abcam | ab219800 |
| Rabbit anti-pRIP3 (Thr231/Ser232) | Cell Signaling Technology | 91702 |
| Mouse anti-β-actin | Sigma | A1978 |
| Rabbit anti-cleaved PARP | Cell Signaling Technology | 9548 |
| Rabbit anti-cleaved caspase-3 (Asp175) | Cell Signaling Technology | 9661 |
| Rat anti-F4/80 | BioLegend | 123101 |
| Rabbit anti-F4/80 | Cell Signaling Technology | 70076 |
| Rabbit anti-CD45 | Cell Signaling Technology | 70257 |
| Goat anti-CD206 | ThermoFisher Scientific | PA5-46994 |
| Rabbit anti-CD8 | Bioss | 0648R |
| Rabbit anti-CD31 | Cell Signaling Technology | 77699 |
| Rabbit anti-pSTAT1 (Y701) | Cell Signaling Technology | 9167 |
| Rat anti-Ki-67 | ThermoFisher Scientific | 14-5698-82 |
| Rabbit anti-SOX10 | abcam | ab180862 |
| Donkey anti-Rat Alexa Fluor 488 | ThermoFisher Scientific | A-21208 |
| Donkey anti-Rabbit Alexa Fluor 555 | ThermoFisher Scientific | A-31572 |
| Donkey anti-Goat Alexa Fluor 647 | ThermoFisher Scientific | A-21447 |
| Rat anti-F4/80 (PE) | BioLegend | 123110 |
| Rat anti-MHC-II (BV650) | BioLegend | 107641 |
| Rat anti-B220 (APC) | ThermoFisher Scientific | 17-0452-82 |
| Rat anti-Ly6C (APC-Cy7) | ThermoFisher Scientific | 560596 |
| Rat anti-CD11b (eFluor 450) | ThermoFisher Scientific | 48-0112-82 |
| Rat anti-CD115 (PE-dazzle 594) | BioLegend | 135527 |
| Rat anti-CD45 (Alexa Fluor 488) | BioLegend | 109816 |
| Rat anti-CD11c (PE/Cy7) | ThermoFisher Scientific | 25-0114-81 |
| TruStain FcX | BioLegend | 101320 |
| IRDye® 680RD Goat anti-Mouse IgG (H + L) | LI-COR | 926-68070 |
| IRDye® 800CW Goat anti-Rabbit IgG | LI-COR | 926-32211 |
| Chemicals and recombinant Proteins |  |  |
| DMEM, high glucose, pyruvate, no glutamine | ThermoFisher Scientific | 10313021 |
| L-Glutamine (200 mM) | ThermoFisher Scientific | 25030081 |
| Penicillin-Streptomycin-Glutamine (100X) | ThermoFisher Scientific | 10378016 |
| Endothelial Cell Growth Medium 2 | PromoCell | C-22011 |
| DPBS, no calcium, no magnesium | ThermoFisher Scientific | 14190144 |
| Dimethyl sulfoxide | Sigma | D4540 |
| Distilled Water | ThermoFisher Scientific | 15230162 |
| Fetal Bovine Serum, certified, heat inactivated | ThermoFisher Scientific | 10082147 |
| Fetal Bovine Serum, qualified, heat inactivated | ThermoFisher Scientific | 16140071 |
| Ethylenediaminetetraacetic acid disodium salt solution | Sigma | E7889 |
| Polyethylene glycol | Sigma | 202371 |
| 10X Tris/CAPS | Bio-Rad | 1610778 |
| TEMED | Sigma | E4378 |
| Triton X-100 | Sigma | X100 |
| Tween 20 | AppliChem | A4974 |
| Trizma Base | Sigma | T1503 |
| Glycine | VWR | 444495D |
| 2-Mercaptoethanol | Merck | 805740 |
| 30% Acrylamide/Bis Solution 37.5:1 | Sigma | A3699 |
| DL-Dithiothreitol | ThermoFisher Scientific | R0861 |
| Sodium Dodecyl Sulfate | Sigma | L3371 |
| Histofix | Roth | P087 |
| Neo-Clear | Merck | 1.09843 |
| Hydrogen peroxide solution | Sigma | H1009 |
| Streptavidin-HRP | Leica | RE7104-CE |
| AEC+ High Sensitivity Substrate Chromogen | Agilent | K3461 |
| Aquatex | Merck | 108562 |
| 10X RIPA Buffer | abcam | ab156034 |
| 4X Protein Sample Loading Buffer | LI-COR | 928-40004 |
| Intercept® (TBS) Blocking Buffer | LI-COR | 927-60001 |
| Intercept® T20 (TBS) Antibody Diluent | LI-COR | 927-65001 |
| Odyssey® Nitrocellulose Membranes | LI-COR | 926-31092 |
| FITC Annexin V | BioLegend | 640906 |
| 7-Aminoactinomycin D | Sigma | A9400 |
| BIX 01294 | Selleckchem | S8006 |
| SKI-II | Selleckchem | S7116 |
| Masitinib | Selleckchem | S1064 |
| Tacedinaline | Selleckchem | S2818 |
| AZD4547 | Selleckchem | S2801 |
| Prochlorperazine dimaleate salt | Selleckchem | S4631 |
| LRRK2-IN-1 | Selleckchem | S7584 |
| Tubastatin A TFA | Selleckchem | S0709 |
| GSK-J4 HCl | Selleckchem | S7070 |
| XMD8-92 | Selleckchem | S7525 |
| Ibrutinib | Selleckchem | S2680 |
| Vandetanib | Selleckchem | S1046 |
| Quizartinib | Selleckchem | S1526 |
| GSK126 | Selleckchem | S7061 |
| PFI-1 | Selleckchem | S1216 |
| Nigericin sodium salt | Selleckchem | S6653 |
| Z-VAD-FMK | Selleckchem | S7023 |
| BV-6 | Selleckchem | S7597 |
| EC359 | MCE | HY-120142 |
| ODN-1826 | InvivoGen | tlrl-1826 |
| Nigericin sodium salt | Sigma | SML-1779 |
| Recombinant Murine M-CSF | PeproTech | 315-02 |
| Animal-Free Recombinant Murine TNF-α | PeproTech | AF-315-01A |
| Aprotinin | Sigma | A1153 |
| Leupeptin | Sigma | L2884 |
| Trypsin Inhibitor | Sigma | T9003 |
| AEBSF | Sigma | A8456 |

#### Critical Commercial Assays

|  |  |  |
| --- | --- | --- |
| PrestoBlue™ Cell Viability Reagent | ThermoFisher Scientific | A13262 |
| CyQUANT™ LDH Cytotoxicity Assay | ThermoFisher Scientific | C20300 |
| CellTiter-Glo® Luminescent Cell Viability Assay | Promega | G7572 |
| Pierce™ Rapid Gold BCA Protein Assay Kit | ThermoFisher Scientific | A53225 |
| Click-iT™ Plus TUNEL Assay | ThermoFisher Scientific | C10619 |
| Streptavidin/Biotin Blocking Kit | VectorLabs | SP-2002 |
| Vector® TrueVIEW® | VectorLabs | SP-8400-15 |

#### Software

|  |  |
| --- | --- |
| GraphPad Prism (Version 9.0.2) | Dotmatics |
| ImageJ Fiji (Version 2.5.0) | ImageJ |
| HALO Image Analysis Software (Version 3.4) | Indica Labs |

Supplementary Figures

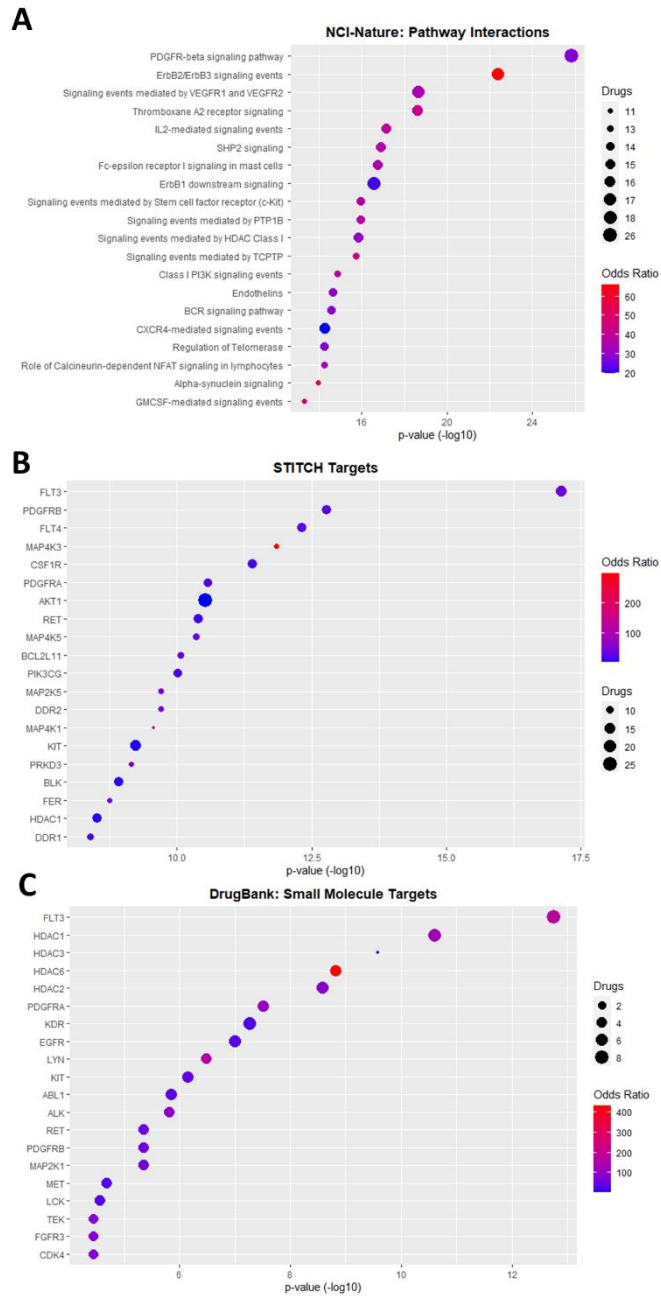

**Figure S1: Bioinformatic analyses of compounds with macrophage-killing activity. (A)** Predicted drug pathway interactions based on pathways contained in the NCI-Nature database. **(B)** Known drug-protein targets present in the EMBL STITCH database. **(C)** Analysis of targeted genes by small molecules contained in the DrugBank database.

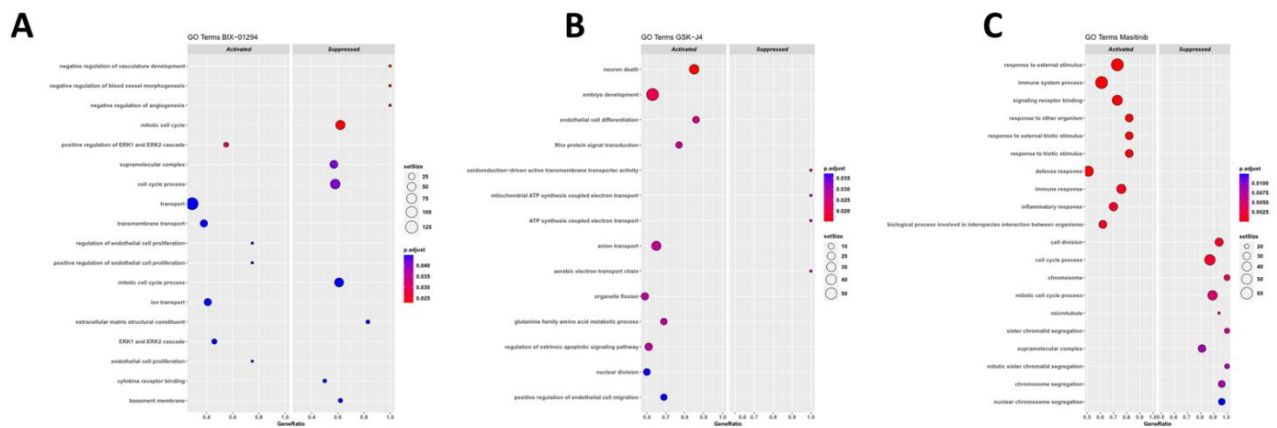

**Figure S2: RNA-sequencing analysis of BMDMs treated with BIX-01294, GSK-J4, or Masitinib. (A – C) Pathway analysis of top up- and downregulated pathways upon treatment with BIX-01294 (A), GSJ-J4 (B), and Masitinib (C). (D) Venn-Diagram depicting differentially expressed genes (DEG) for the individual drug treatments and overlaps of DEG for multiple drugs.**

A

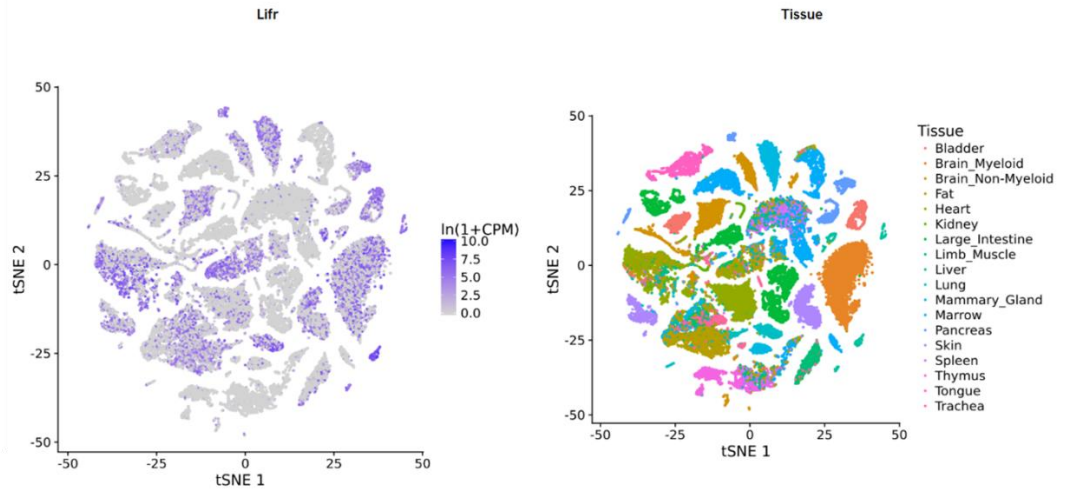

B

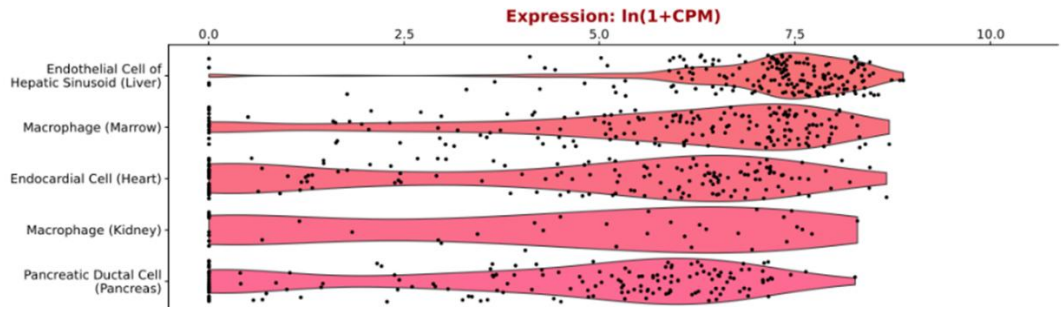

**Figure S3:** (A) Gene expression of *Lifr* (left panel) in the Tabula muris database. The right panel shows the annotations for specific tissues. (B) Cell type specific expression of *Lifr* of the 5 cell types with the highest expression in Tabula muris database.

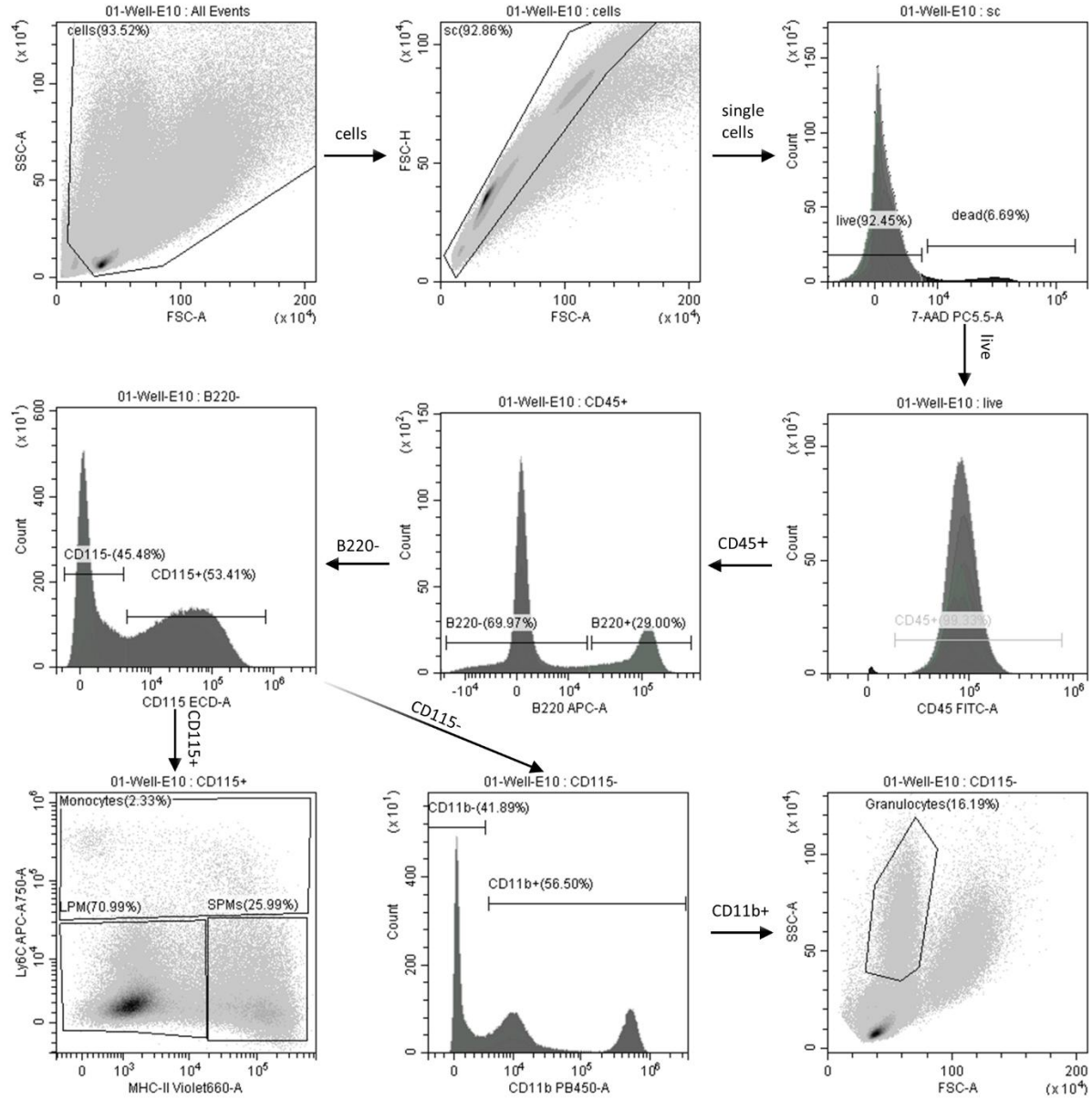

**Figure S4:** Gating strategy for flow cytometric analysis of peritoneal cells.

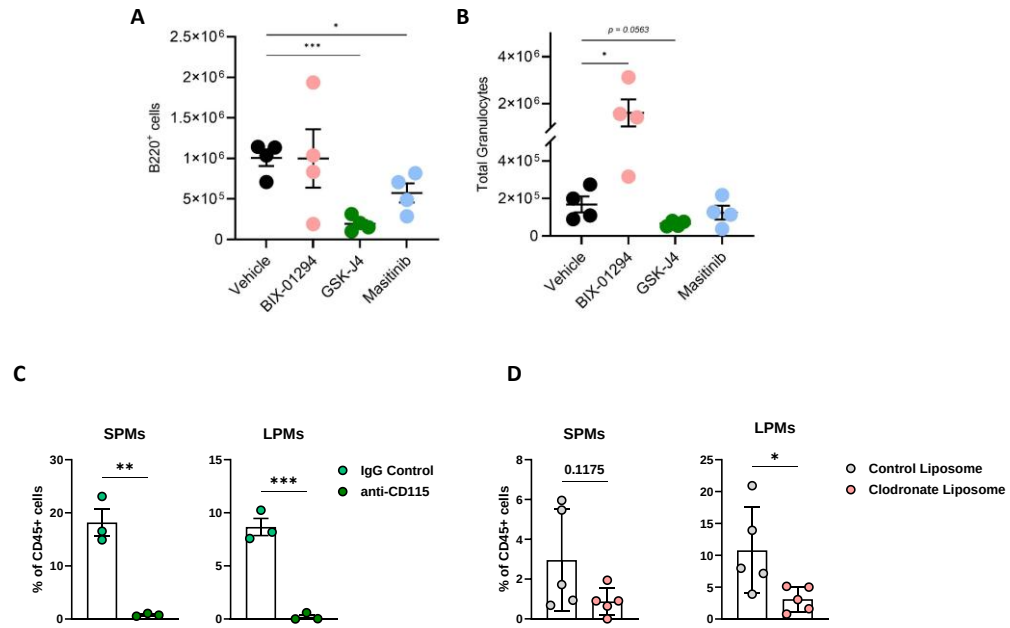

**Figure S5:** Quantification of B-cells (B220+ cells; **A**) and granulocytes (**B**) in peritoneal exudate in accordance to Figure 4. Quantification of small (SPM) and large peritoneal macrophages (LPM) upon macrophage depletion with anti-CD115 antibody (**C**) or clodronate liposomes (**D**).

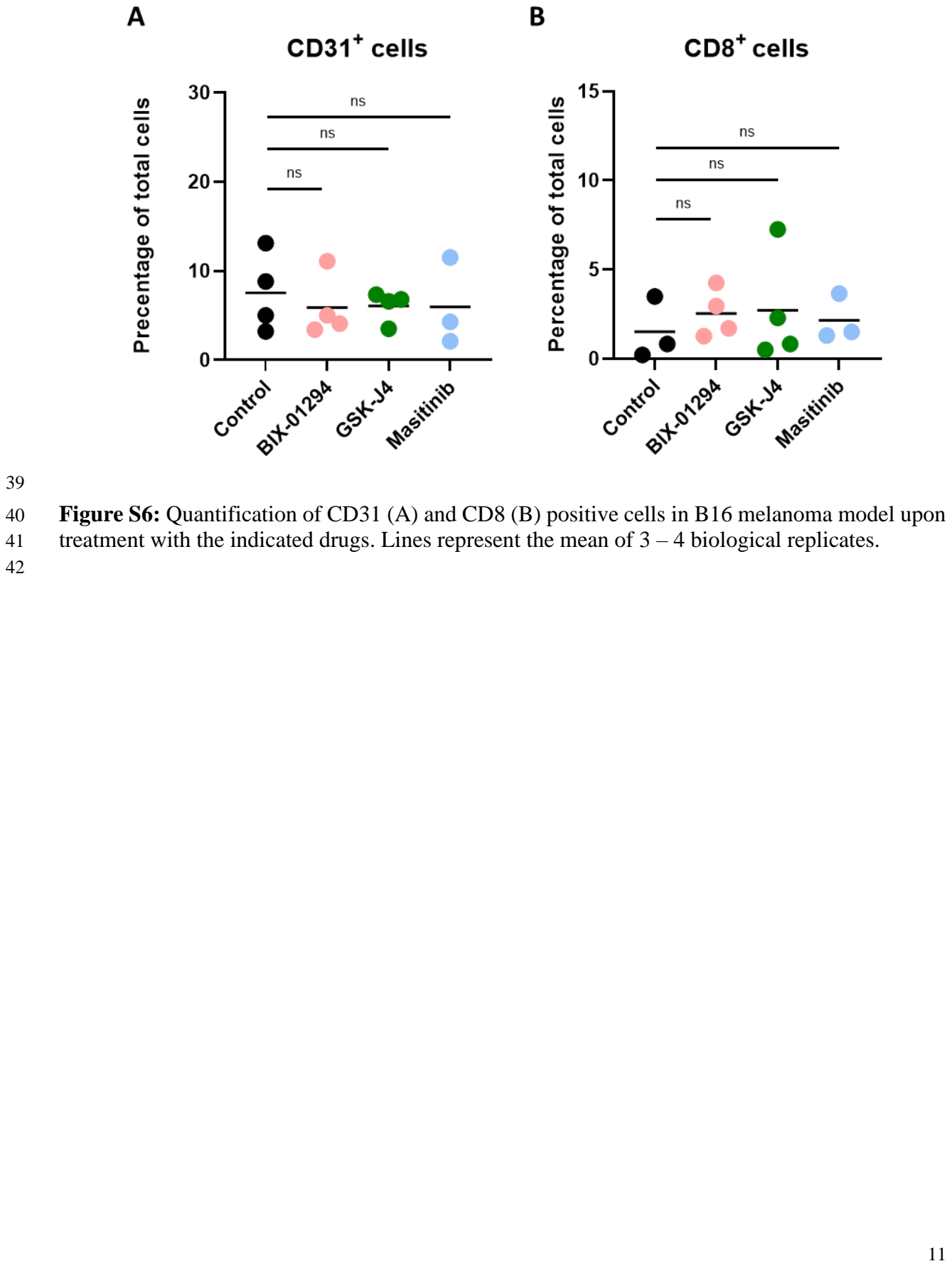

Figure S6: Quantification of CD31 (A) and CD8 (B) positive cells in B16 melanoma model upon treatment with the indicated drugs. Lines represent the mean of 3 – 4 biological replicates.

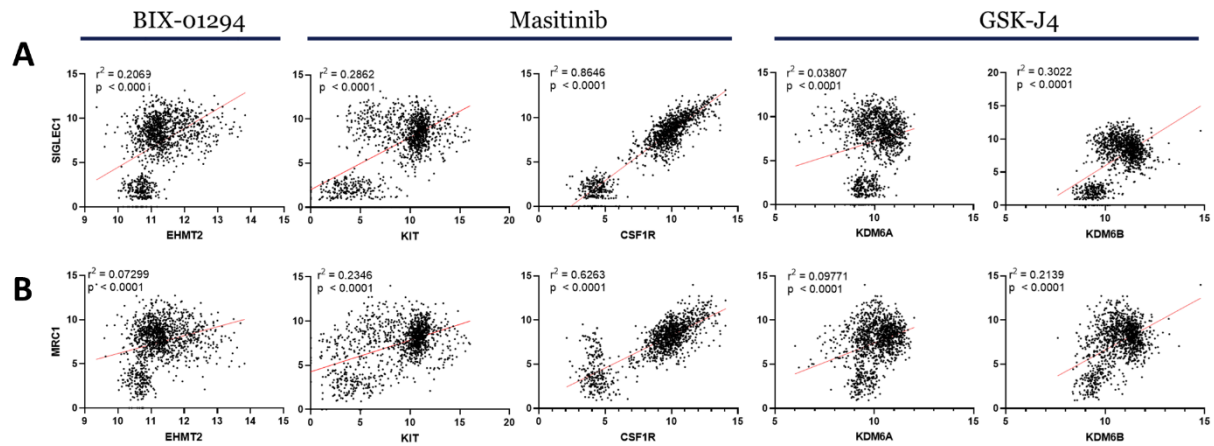

**Figure S7:** Correlation between target genes of BIX-01294 (EHMT2), Masitinib (KIT, CSF1R), and GSK-J4 (KDM6A, B) and TAM genes SIGLEC1 (A) and MRC1 (B).
